# Spreading α-synuclein rewires organelle communication and impairs neuron–astrocyte mitochondrial quality control

**DOI:** 10.64898/2026.06.19.733342

**Authors:** Elisabeth Fritsch, Kamelija Horvatovic, Paula Santos Otte, Tomas Koudelka, Jana Rossius, Caroline Braeuning, Laura Breimann, Ilaria Piazza, Melissa Birol

**Author notes:** Correspondence should be addressed to: Melissa Birol. Equal contribution (Kamelija Horvatovic, Paula Santos Otte).

## Abstract

Progressive intercellular spreading of α-synuclein (αS) is implicated in pathology initiation and propagation of synucleinopathies. However, how recipient neurons respond to incoming αS and whether these responses contribute to disease-associated early metabolic events, remains unknown. Here, using extracellular monomeric αS to model the earliest cellular response to spreading, we found that internalized αS accumulates at tri-organelle contact sites linking mitochondria, endoplasmic reticulum, and endo/lysosomal compartments. At these interfaces, αS stabilizes generally dynamic contacts and constrains their remodeling, thereby rewiring organelle communication. These effects require the acidic αS C-terminus and are not recapitulated by intracellular αS overexpression. Proteomic profiling of αS-associated mitochondria identified a contact site-enriched but quality-control-deficient state. Functionally, spreading αS impairs neuron-astrocyte mitochondrial quality control (MQC) by reducing neuronal mitochondria transfer to astrocytes, while enhancing mitochondrial import. Our findings establish organelle contact sites as critical target of spreading αS, through which rewired organelle communication impairs MQC and neuron-astrocyte crosstalk.

## Introduction

The propagation of pathogenic proteins between cells is increasingly recognized as a fundamental process in neurodegenerative diseases^1,2^. In synucleinopathies, α-synuclein (αS) spreads through interconnected neural circuits and is thought to contribute to the initiation and progression of pathology^3–5^. Both monomeric and pathological forms of αS spread between cells^4,6,7^, exploiting routes such as tunneling nanotubes, exosomes, and receptor-mediated endocytic uptake to move from neuron to neuron or from neurons to glial cells^8–12^. Consistent with this αS is detected in cerebrospinal fluid (CSF) under both physiological and pathological conditions^13,14^, indicating that neurons continuously release αS into the extracellular space. Such secretion may either confer specific biological functions or serve as a clearance route to prevent aggregation, whereas reuptake by neighboring cells may drive disease progression. Basal extracellular αS levels in CSF are typically reported in the low nanogram-per-milliliter range, although absolute concentrations vary across studies and detection assays used^14,15^. In synucleinopathies, the total CSF αS is typically reduced, whereas oligomeric and phosphorylated species increase^16^, changes thought to reflect the sequestration of αS into intracellular aggregates. Once internalized, oligomeric and fibrillar αS species can overwhelm intracellular homeostasis, particularly the autophagy-lysosome system, leading to further accumulation and cellular stress^17,18^. In parallel, glial cells such as astrocytes and microglia respond to extracellular αS by releasing pro-inflammatory cytokines that aggravate neuronal dysfunction^19–21^. While these αS species are established drivers of proteostatic stress, much less is known about the effects of monomeric αS that is continuously exchanged between cells under physiological and pathological conditions. Whether monomeric αS transfer alone can reshape cellular organization and homeostasis is unknown. Addressing this question is essential for defining the earliest cellular consequences of αS spreading and identifying mechanisms that may predispose recipient cells to later dysfunction.

Mitochondrial dysfunction is among the earliest and most prominent features of synucleinopathies^22–25^. In these disorders, αS impairs mitochondrial function^26^ and co-accumulates with mitochondria within Lewy bodies^27^. Disruption of mitochondrial dynamics leads to energy deficits, oxidative stress, and excessive production of reactive oxygen species (ROS), thereby accelerating neuronal degeneration^26,28–30^. Beyond intrinsic defects, mitochondrial function depends on coordinated communication with other organelles, including the endoplasmic reticulum (ER)^31–33^ and the endolysosomal system^34,35^. These interactions are mediated through specialized membrane contact sites, including mitochondria-associated membranes (MAMs), which integrate lipid transfer, calcium signaling, and mitochondrial dynamics and are increasingly recognized as key regulators of mitochondrial quality control (MQC)^36–38^. Moreover, controlled cell-to-cell communication has recently been shown to be crucial for MQC^39–41^, highlighting the interplay between intra- and intercellular mechanisms that sustain mitochondrial function. Notably, αS has been reported to associate with MAMs and to modulate ER–mitochondria communication^42–45^. In Parkinson’s disease (PD), both rare familial mutations and common genetic risk variants converge on the endolysosomal system and mitochondria as key mediators of cellular dysfunction and neurodegeneration. Genes such as LRRK2, GBA1, VPS35, and ATP13A2 regulate endolysosomal homeostasis, whereas PINK1 and PRKN control mitophagy^46,47^.

Mitochondria themselves can be exchanged between cells, a process known as mitochondrial transfer^41,48^. Damaged or dysfunctional mitochondria can be replaced by intact mitochondria from healthier cells, promoting cell survival and function^41,49,50^. Stressed cells can extrude damaged mitochondria that are then taken up and degraded by recipient cells^39,40^. Mitochondrial transfer allows cells to share bioenergetic capacity, compensate for impaired ATP production, mitigate oxidative stress, and restore mitochondrial function in compromised cells. This process has been observed in tissue repair^50^, immune responses^51^, and diseases such as cancer^52,53^, cardiovascular disorders^40^, and, more recently, in neurodegeneration^54,55^. In the brain, glial cells play a key supportive role in maintaining neuronal health and homeostasis^56^. Under mitochondrial stress, neurons can extrude dysfunctional mitochondria via tunneling nanotubes or vesicular pathways, and neighboring astrocytes can engulf these mitochondria and degrade them through their own lysosomal machinery^39^. This process depends on coordinated mitochondrial dynamics, vesicular trafficking, and organelle contact-site signaling. This intercellular MQC mechanism enables neurons to offload compromised mitochondria, preventing the accumulation of ROS and progressive mitochondrial failure. Whether the spreading of αS interferes with these mechanisms remains unknown.

To determine whether spreading αS interferes with intracellular and intercellular MQC mechanisms, we investigated how recipient neurons respond to incoming αS. Using extracellular monomeric αS to model early spreading-associated uptake, we found that internalized αS accumulates at organelle contact sites linking mitochondria, the ER, and the endolysosomal system, where it stabilizes normally dynamic MAM-endosome contacts and constrains their remodeling. Quantitative imaging, analyses of organelle dynamics, and proteomic profiling further revealed that αS selectively remodels contact site-associated mitochondrial subpopulations, promoting a contact site-enriched but quality control-deficient mitochondrial state characterized by altered organelle communication and impaired MQC. Functionally, these changes compromise neuron–astrocyte MQC by reducing the transfer of damaged neuronal mitochondria to astrocytes while enhancing compensatory mitochondrial delivery from astrocytes to neurons under stress conditions. Together, our findings establish a mechanistic link between αS spreading, organelle communication, and intercellular mitochondrial homeostasis, providing insight into how early αS transfer initiates cellular dysfunction in synucleinopathies.

## Results

### Spreading αS localizes on mitochondria-ER-endo/lysosomal contacts

The spread of αS across neurons is thought to involve its trafficking through interconnected organellar networks, yet how internalized αS engages specific subcellular compartments remains unclear. To define αS intracellular localization upon spreading, iPSC-derived dopaminergic neurons (DNeurons) and SH-SY5Y cells were conditioned with monomeric ATTO 700-labeled αS, at concentrations of 80 nM (denoted αS) and 250 nM (denoted αS_D_) for 14 h, unless otherwise indicated. These concentrations were selected to assess dose-dependent effects on intracellular targeting while remaining non-toxic to neurons over the experimental time course^11^. Although higher than αS levels typically detected in CSF^14,15^, these concentrations were used to model localized exposure in recipient cells upon intercellular transfer. Following internalization, αS accumulated as discrete puncta that localized predominantly at the termini of mitochondrial tubules (Fig. 1a and S1a). Line-profile analyses confirmed that αS puncta resided adjacent to, but rarely overlapped with, mitochondrial tubules (Fig. 1b and S1b). We next performed fluorescence-activated organelle sorting (FAOS) of cell lysates obtained from cells conditioned with αS or αS_D_ (Fig. 1c). Using MitoTracker Green FM and Alexa Fluor 647-labeled αS, we resolved and quantified distinct mitochondrial subpopulations co-sorting with αS and αS_D_. This revealed a significant dose-dependent increase in mitochondrial association, with approximately 3.93 ± 0.99% and 7.47 ± 0.50% of total mitochondria events detected for αS and αS_D_, respectively (Fig. 1c, S1c-e). These data demonstrate that a defined subset of mitochondria associates with internalized αS.

**Figure 1.**
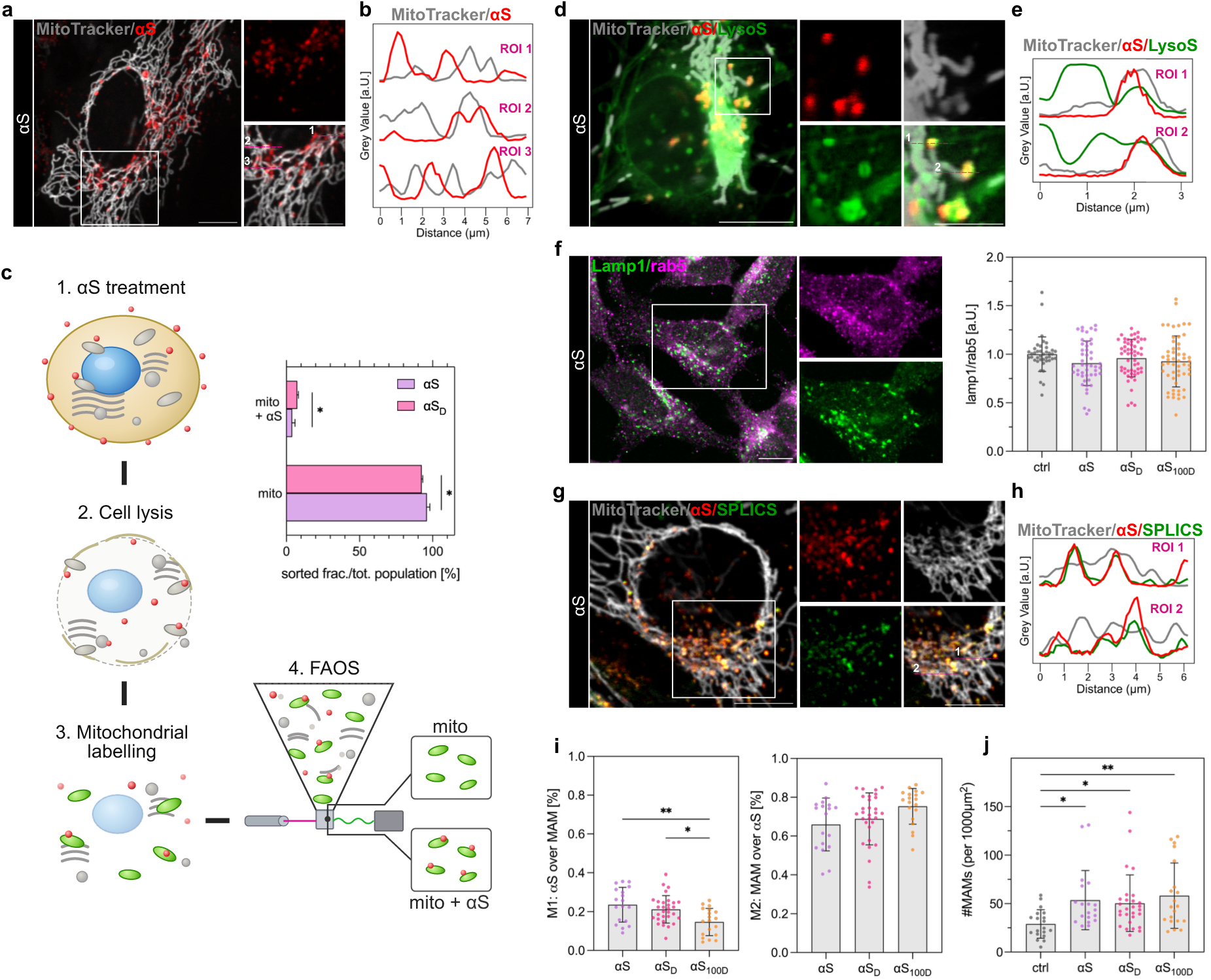
Internalized monomeric αS localizes to mitochondria and tri-organelle contact sites. a, Representative confocal images of SH-SY5Y cells labeled with MitoTracker (grey) following treatment with αS (red). Insets highlight regions of mitochondria-associated αS puncta. b, Fluorescence intensity line profiles across selected regions of interest (ROIs) from (a) showing αS puncta in close apposition to, but only partial overlap with mitochondria. c, Schematic of FAOS workflow for isolation of mitochondria. SH-SY5Y were treated with αS, lysed, and MitoTracker-labeled mitochondria were sorted from whole cell-lysates into αS-positive (αS⁺) and αS-negative (αS⁻) sub-populations for downstream analysis. Fractions of αS⁺ and αS⁻ mitochondria of total sorted population shown in quantification. Unpaired two-tailed t-test with Welch’s correction. *N=2-4* biological replicates; total sorted single events ranged from ∼36000 to 290000. d, Confocal images of dopaminergic neurons labeled with MitoTracker, αS, and LysoSensor (green), revealing frequent localization of mitochondria-associated αS puncta with endolysosomal compartments. Insets highlight regions containing mitochondria, αS and endolysosomes. e, Intensity line profiles of ROIs from (d) demonstrating overlap between αS and LysoSensor-positive compartments adjacent to mitochondria. f, Immunofluorescence images of SH-SY5Y co-stained for late endosomal-lysosomal (Lamp1) and early endosomal (Rab5) markers. Lamp1/Rab5 fluorescence intensity ratio is unchanged across all conditions indicating that αS treatment does not affect endolysosomal pool composition. Kruskal-Wallis with Dunn’s post-hoc. *N=*5-7 replicates. g, Representative images of SH-SY5Y expressing MAM-marker SPLICS (green), together with MitoTracker and αS. Insets highlight αS-MAM localization. h, Intensity line profiles of ROIs from (g) demonstrating colocalization of αS with SPLICS-positive contact sites. i, Object-based colocalization analysis of αS with SPLICS. M1 (fraction: αS overlapping MAM) was significantly reduced for αS_100D_. M2 (fraction: MAM overlapping αS) showed no significant difference across αS variants. Ordinary one-way-Anova with post hoc Tukey’s. *N=*2-3 biological replicates. j, Density of SPLICS-positive MAM contact sites per 1000 µm² cell area. αS treatment increased MAM density compared to untreated control, with no significant differences detected between αS variants. Kruskal-Wallis with Dunn’s post-hoc. *N=2-3* biological replicates. Data are presented as mean ± s.e.m: each point represents a single FOV. * p < 0.05, ** p < 0.01). Scale bars: a, d, f, g, 10 µm; d inset, 5 µm.

Consistent with this, mitochondria-associated αS puncta frequently appeared within endosomal compartments, suggesting receptor-mediated endocytosis as the principal uptake pathway^10,11^. We found that mitochondria-adjacent αS puncta were often detected at endosomal compartments in SH-SY5Y cells and in DNeurons (Fig. 1d and S1f). Line intensity profiles revealed substantial overlap between αS and LysoSensor-positive endo/lysosomal compartments that reside in proximity to mitochondria (Fig. 1e and S1g). Given that the acidic C-terminal domain of αS regulates membrane interactions, we next tested its contribution to endolysosomal association using a truncated variant comprising residues 1-100 (denoted αS_100D_), applied at 250 nM to match the αS_D_ condition. To determine whether αS preferentially associates with specific endolysosomal subpopulations or alters organelle acidification, we quantified LysoSensor fluorescence intensity at the level of individual compartments. LysoSensor intensity was used to assess endolysosomal acidity, enabling comparison between the total endolysosomal population (Fig. S1h) and the subset associated with αS (Fig. S1i). αS-localized compartments were modestly enriched within the more acidic range, however, the overall distribution was not shifted across conditions (Fig. S1h, i). While αS_D_ displayed a broader distribution and αS_100D_ a more constrained profile, the overall tendency remained unchanged. To assess whether αS spreading alters the composition of the endolysosomal pool, we quantified early (Rab5) and late (Lamp1) endosomal markers. Their relative abundance remained unchanged, indicating that αS does not shift the balance between early and late compartments (Fig. 1f).

Endosomes communicate with mitochondria through ER-defined contact sites known as MAMs^57,58^, and notably, αS has been reported to associate with MAMs^42–45^. Together, this raises the possibility that spreading αS accumulates at interfaces coordinating endolysosomal–mitochondrial crosstalk. To test this, SH-SY5Y cells expressing the split-GFP-based MAM contact-site reporters (SPLICS_S_-P2A^ER–MT^, to detect interactions over the range of ∼8-10 nm^59^) were treated with αS and imaged. Internalized αS localized to SPLICS-positive puncta that overlapped with the MAM tether protein VAPB (Fig. 1g, h and S1j-m). Colocalization across αS-treated conditions was confirmed by fluorescence intensity line profiles and object-based colocalization analysis (Fig. 1h, i and S1k, m, n). Measurement of the fraction of total αS overlapping MAMs (M1) revealed comparable enrichment for αS and αS_D_, with approximately 20% of total αS puncta associated with MAMs. In contrast, αS_100D_ exhibited a reduction in MAM-associated αS signal (Fig. 1i and S1n). When normalized to total MAMs (M2), all αS variants occupied a substantial fraction (>60%) of available MAM structures. These data suggest that αS engages MAMs as part of its intracellular trafficking route, with the C-terminus modulating its relative enrichment at these interfaces. We next asked whether αS association alters MAM architecture. Quantification of contact-site density revealed an increase in MAM number across all αS conditions compared with control cells (Fig. 1j). Although mean MAM size was unchanged (Fig. S1o), size-distribution analysis revealed a shift toward smaller contact areas, most prominently in αS_D_. These findings demonstrate that αS not only localizes to MAMs but also actively remodels contact-site architecture, positioning these interfaces as key sites of αS-mediated organelle regulation.

### Spreading αS alters the dynamic remodeling of MAM-endosome contacts

Organelle contact sites occupy a small fraction of total membrane surface yet play essential roles in lipid, ion, and metabolite exchange^60^. These contacts are highly transient, continuously forming and resolving to maintain organelle communication. We hypothesized that αS recruitment to MAM-endosome interfaces may impair this dynamic remodeling process, thereby restricting organelle plasticity. To probe these dynamics, we employed a hypotonic swelling assay that induces the formation of large intracellular vesicles (LICVs), expanding organelle membranes and spatially separating compartments while preserving inter-organelle contacts. This reorganization enables real-time visualization of membrane remodeling and contact-site dynamics that are otherwise unresolved in intact cells^61^.

SH-SY5Y cells and DNeurons were stained with MitoTracker Red CMXRos to visualize intact mitochondria. Under isotonic conditions, mitochondria in both control and αS-conditioned cells displayed a typical branched tubular network (Fig. S2a), with a modest reduction in network complexity observed for αS_100D_. Upon hypotonic treatment, these networks rapidly fragmented into numerous mitochondrial LICVs over 30 min, reflecting normal remodeling between mitochondria (Fig. 2a and MovieS1). This assay thus provides a sensitive readout for contact-site flexibility and mitochondrial fission-fusion dynamics. The extent and kinetics of mitochondrial fragmentation and LICV formation reflect the capacity for membrane remodeling at ER-mitochondria interfaces. We next examined how the presence of spreading αS affects this process. DNeurons and SH-SY5Y cells were pre-exposed to αS for 14 h before undergoing hypotonic swelling, and organelle remodeling was monitored by time-lapse imaging (Fig. 2a and MovieS2 (αS), MovieS3 (αS_D_)). Under hypotonic conditions, control cells showed extensive vesiculation. In contrast, αS-treated cells retained elongated mitochondrial structures. Quantitative segmentation of these images at the end of hypotonic treatment (30 min), revealed that the presence of αS markedly reduced LICV number while preserving elongated mitochondria (Fig. 2b-e). Mitochondria showed a dose-dependent decrease in LICV number following αS exposure (Fig. 2b, c), accompanied by increased mitochondrial branch length (Fig. 2d) and aspect ratio, a measure of length-to-width proportion (Fig. 2e). These concentration-dependent phenotypic differences were comparable between SH-SY5Y and DNeurons (Fig. S2b-d). Similar to full-length αS_D_, αS_100D_ reduced mitochondrial LICV number; however, mitochondrial morphology, including branch length and aspect ratio, remained comparable to that of untreated controls (Fig. 2c-e, S2b-e, and MovieS4 (αS_100D_)). These findings indicate that αS restricts mitochondrial fragmentation and reshaping in response to cellular stress, consistent with impaired fission-fusion activity at these organelle contact zones.

**Figure 2.**
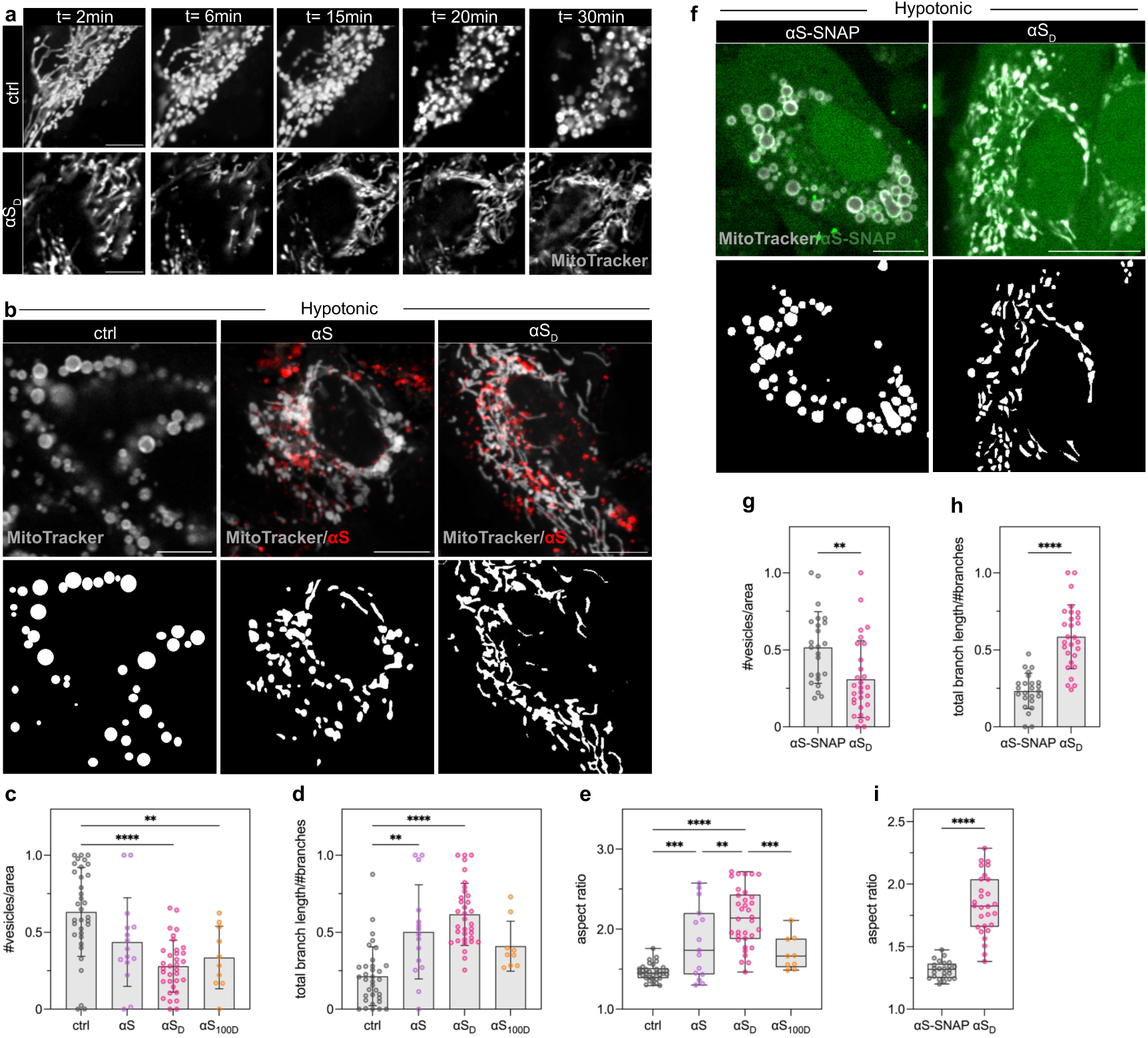
Spreading αS constrains mitochondrial remodeling and organelle plasticity. a, Time-lapse confocal imaging of mitochondria after hypotonic treatment in ctrl and αS_D_-treated SH-SY5Y cells. Representative frames show mitochondrial network dynamics over time (2–30 min post-hypotonic treatment), revealing reduced mitochondrial remodeling and motility in αS_D_. b, Representative images of mitochondria ∼30 min post-hypotonic swelling, visualized by MitoTracker in ctrl and αS_D_ cells. Bottom panels show corresponding segmentation of mitochondria. c-e, Quantifications of mitochondria morphology after hypotonic swelling, comparing the number of LICVs per cell area (c), mean branch length (d), and mitochondria tubule elongation (aspect ratio, e), between ctrl and αS conditions. αS exposure impairs mitochondrial vesiculation, in a concentration- and C-terminal-dependent manner. Ordinary one-way-Anova with Tukey’s post-hoc. *N=*2-5 biological replicates. f, Representative images of LICVs in αS-SNAP overexpressing SH-SY5Y cells under hypotonic conditions. Conditioning with αS_D_ shows impaired mitochondrial vesiculation. g-i, Quantification of mitochondria morphology post-hypotonic treatment of αS-SNAP overexpressing SH-SY5Y cells, conditioned with- (denoted αS_D_) and without-(denoted αS-SNAP) αS_D_. Unpaired two-tailed Student’s t-test or Mann-Whitney. *N=*2 biological replicates. Data are presented as mean ± s.e.m.; each point represents a single FOV. * p < 0.05, ** p < 0.01, *** p < 0.001, **** p < 0.0001. Scale bars: 10 µm.

Finally, to distinguish whether these effects arise from extracellular spreading αS or increased intracellular αS abundance, SH-SY5Y cells overexpressing αS-SNAP were analyzed. Although total αS levels were increased several-fold relative to cells treated with spreading αS (Fig. S2f), overexpression alone did not affect LICV formation, mitochondrial branch length, or aspect ratio (Fig. 2f-i and S2g). In contrast, when cells expressing αS-SNAP that were also conditioned with extracellular αS, this recapitulated the characteristic phenotypes resulting in reduced vesiculation, and elongated mitochondria (Fig. 2f-i). These data demonstrate that specifically spreading αS recruited to inter-organelle contacts, rather than intracellular αS accumulation, disrupts the dynamic remodeling of MAM-endosome interfaces and constrains mitochondrial fission-fusion behavior. Together, these results indicate that αS accumulation at contact sites limits organelle membrane plasticity, likely by stabilizing otherwise transient MAM-endosome junctions.

### Spreading αS reprograms mitochondria into a contact site-enriched, quality control-deficient state

Having established that spreading αS localizes to mitochondria-ER-endolysosomal contact sites, remodels MAM architecture, and constrains mitochondrial dynamics, we next sought to define the molecular machinery selectively associated with αS-engaged mitochondria. We reasoned that identifying this protein environment would elucidate the interorganelle pathways most directly perturbed by αS and provide mechanistic insight into downstream functional consequences. To this end, we performed quantitative LC–MS/MS-based proteomics on mitochondria isolated from control and αS_D_-conditioned cells. Mitochondria were FAOS-sorted based on MitoTracker and internalized αS_A647_ signal. This approach yielded three distinct populations: control mitochondria (ctrl) from untreated cells, mitochondria co-sorted with internalized αS (αS⁺ mitochondria), and mitochondria from αS-treated cells lacking detectable αS signal (αS⁻ mitochondria, Fig. 3a). Importantly, αS⁻ mitochondria represent an internal matched control derived from the same αS-exposed cells as αS^+^. This strategy enabled discrimination between the global effects of αS exposure and protein networks specifically associated with αS-engaged mitochondrial populations. Quantitative mass spectrometry revealed that αS⁺ mitochondria exhibited a proteomic profile that was clearly distinct from both ctrl and αS⁻ fractions. Principal component analysis demonstrated robust segregation of αS⁺ mitochondria, while αS⁻ mitochondria clustered closely with ctrl samples (Fig. 3a). This suggested that proteomic remodeling induced by spreading αS is spatially restricted rather than uniform across the mitochondrial network. Differential expression analysis supported this observation. Comparison of αS⁺ mitochondria to ctrl identified significant proteomic changes (2528 over-represented, 1240 under-represented proteins), which were similarly evident when comparing αS⁺ to αS⁻ mitochondria (2601 over-represented, 1180 under-represented proteins). In contrast, αS⁻ mitochondria showed no significant proteomic changes relative to control (Fig. 3b and S3a).

**Figure 3.**
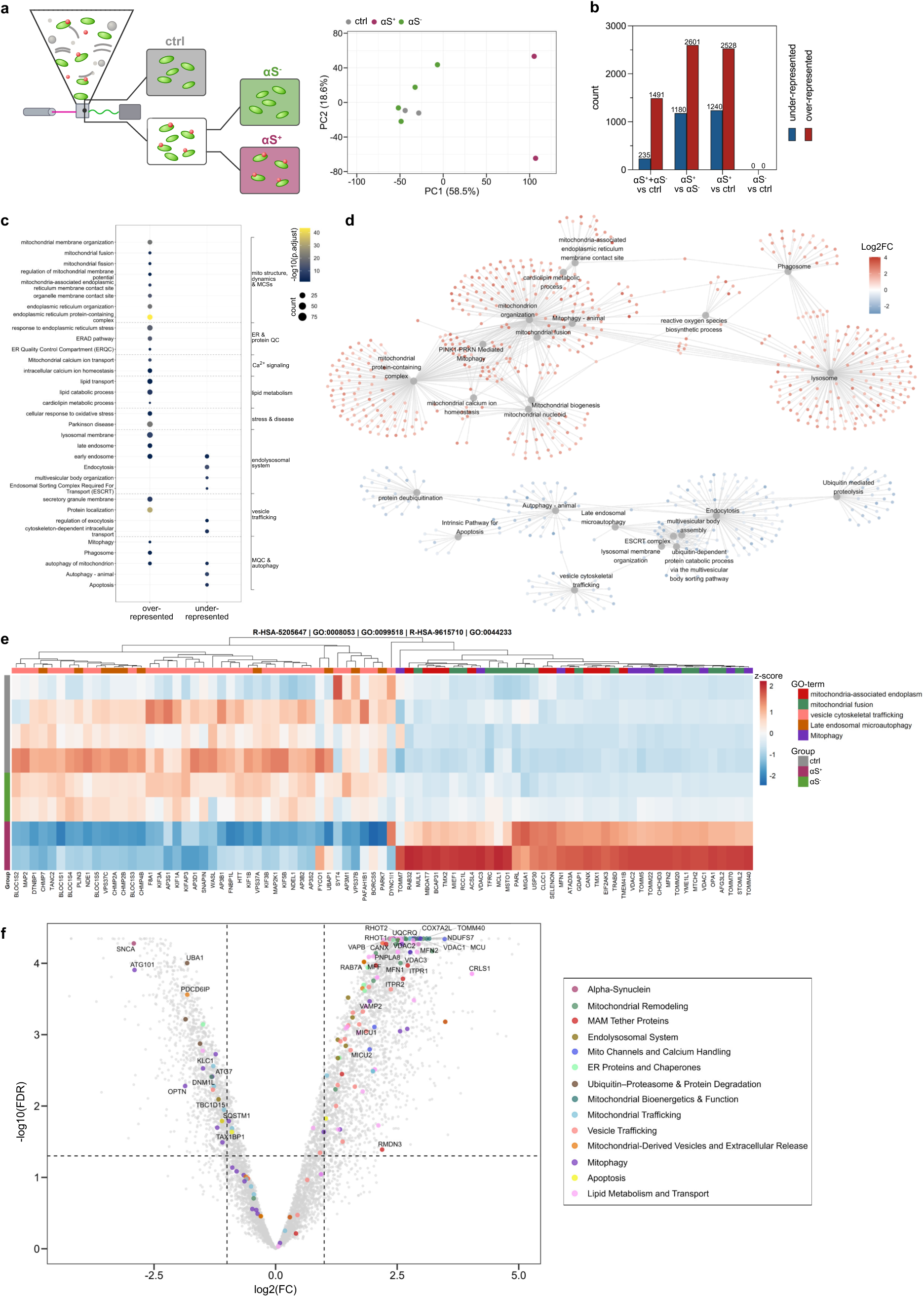
Spreading αS defines a contact-site–enriched mitochondrial proteome with altered metabolic and quality-control pathways. a, FAOS workflow for proteomic profiling of mitochondrial subpopulations. Mitochondria were isolated from control cells (ctrl) and αS-treated cells and separated into αS-associated (αS⁺) and non-associated (αS⁻) fractions. PCA shows a distinct proteomic profile for αS⁺ mitochondria compared with ctrl and αS⁻ fractions. b, Quantification of differentially abundant proteins across comparisons. Bar plot shows the number of over- and under-represented proteins in αS⁺/αS⁻ vs ctrl, αS⁺ vs ctrl, αS⁺ vs αS⁻ and αS⁻ vs ctrl comparisons. αS⁺ mitochondria exhibit extensive proteomic changes relative to ctrl and αS⁻, while αS⁻ vs ctrl shows zero changes. c, ORA dot plot showing pathway terms concordantly over- and under-represented in αS⁺ mitochondria across both αS⁺ vs ctrl and αS⁺ vs αS⁻ comparisons. This identified pathways robustly associated with the αS⁺ proteomic state, consistent with GSEA (Fig. S3b). d, Cnet plot of leading-edge proteins from selected GSEA-enriched pathways (αS⁺ vs ctrl). Proteins associated with mitochondrial dynamics, morphology, and early MQC processes cluster separately from proteins involved in vesicle trafficking, autophagy, and degradative mitophagy pathways, with no overlap between the two networks. e, Heatmap of leading-edge proteins from selected pathway terms shown in the cnet analysis (d) across all conditions (ctrl, αS⁻, αS⁺; rows represent biological replicates). Columns represent individual proteins grouped and color-coded by pathway. Hierarchical clustering confirms the separation of positively and negatively enriched pathways. f, f, Volcano plot of the αS⁺ versus ctrl proteome. Significantly altered proteins are shown in grey, while selected proteins of interest are highlighted and grouped into curated functional categories. Annotated proteins are referenced in the Results section. Dashed lines indicate significance thresholds (adjusted p < 0.05; |log₂FC| > 1). Biological replicates, ctrl, αS, *N=*4; αS_D_, *N=*2.

To define pathways associated with αS-engaged mitochondria, we performed enrichment analysis across the sorted mitochondrial populations. Comparison of combined mitochondrial fractions from αS-treated cells (αS⁺ + αS⁻) against ctrl primarily captured global responses to αS exposure. Pathways associated with mitochondria contact sites, mitochondrial organization, lipid metabolism, and calcium signaling were enriched, consistent with activation of inter-organelle communication networks (Fig. S3b). In contrast, endolysosomal membrane organization and degradative trafficking pathways were depleted. Over-representation analysis (ORA) further confirmed the coherence of these pathways, revealing strong concordance in pathway enrichments between αS⁺ vs ctrl and αS⁺ vs αS⁻ comparisons (Fig. 3c and Fig. S3c). The connectivity network (cnet) representation further organized these pathways into two functionally distinct modules with no leading-edge protein overlap (Fig. 3d). A positively enriched module comprised pathways associated with mitochondrial bioenergetics, membrane architecture, ER–mitochondria contact sites, and mitophagy-related signaling. In contrast, a negatively enriched module contained pathways linked to autophagy execution, vesicle-mediated trafficking, and degradative cargo processing. This organization suggests a functional separation between pathways associated with contact-site engagement and upstream MQC signaling and those required for downstream execution and degradation. Heatmap analysis of leading-edge proteins further supported this functional segregation at the level of individual proteins (Fig. 3e and S3d).

αS⁺ mitochondria were enriched in proteins associated with mitochondrial bioenergetics, membrane organization, and metabolic remodeling (Fig. 3f, Fig. S3e and Supplementary Table 1). These included multiple components of the electron transport chain, such as NDUFS subunits, the Complex III subunit UQCRQ, and the respiratory supercomplex assembly factor COX7A2L, as well as regulators of mitochondrial structure, including OPA1 and TOMM family translocase subunits. Proteins associated with ER–mitochondria tethering and calcium exchange were coordinately enriched, including the MAM-associated tethers VAPB and RMDN3, MFN1/2, VDAC1/2/3, and ITPR1/2, together with components of the MCU, MICU1, MICU2, and LETM. ER-localized MAM proteins, including CANX, BCAP31, and the ER stress kinase EIF2AK3/PERK, were also enriched, supporting activation of ER–mitochondria stress signaling pathways. In parallel, enzymes linked to cardiolipin biosynthesis and membrane remodeling, including CRLS1, DNAJC19, and PNPLA8, were elevated, consistent with reorganization of mitochondrial membrane composition at contact interfaces.

Conversely, proteins associated with downstream mitochondrial turnover and degradative trafficking were coordinately depleted (Fig. 3f, Fig. S3e and Supplementary Table 1). Components of ubiquitin-mediated cargo tagging, including multiple E1/E2/E3 enzymes, were reduced alongside core autophagy machinery (ATG7/12, and ATG13/101) and selective autophagy receptors, including SQSTM1/p62, OPTN, and TAX1BP1. Proteins involved in mitochondrial transport, including kinesin motor subunits (KIF1A and KLC family members), were similarly depleted. In parallel, machinery required for membrane scission, endosomal sorting, and cargo resolution, including CHMP and VPS37 family proteins, PDCD6IP/ALIX, BLOC-1complex subunits, AP-3 adaptor components, and the mitophagy regulator TBC1D15, was reduced, consistent with impaired downstream trafficking and turnover capacity. Notably, endogenous SNCA was also reduced in αS-associated mitochondrial fractions, suggesting redistribution of the native αS pool. Several protein pairs further suggested a selective enrichment of membrane-engagement and docking machinery and the depletion of proteins required for downstream remodeling and trafficking. The mitochondrial fission receptor MFF was enriched, while its effector GTPase DNM1L/DRP1was reduced, consistent with impaired mitochondrial fragmentation in αS-associated mitochondria. Similarly, the trafficking adaptors RHOT1 and RHOT2 were enriched, while kinesin motor proteins were depleted, suggesting preserved mitochondrial docking but reduced transport capacity. Although αS⁻ mitochondria showed no significant protein changes relative to control, ORA identified weak but coordinated pathway-level shifts within this fraction (Fig. S3f). These were primarily linked to RNA-related and mitochondrial stress-response pathways, including mt-tRNA processing, iron–sulfur cluster assembly, and electron transport chain regulation^62^. In contrast, P-body assembly pathways were relatively depleted^63^. Thus, while the major proteomic remodeling was restricted to αS-associated mitochondria, αS exposure also induced lower-magnitude secondary responses. Together, these data suggest that αS engagement promotes an interorganelle contact site-enriched mitochondrial state with reduced association of downstream degradative machinery.

### Spreading αS at tri-organelle contact sites impairs neuron-to-astrocyte mitochondria transfer

The proteomic enrichment of vesicular trafficking and MQC pathways within αS-associated mitochondrial fractions suggested that αS engagement at contact sites may extend beyond intracellular remodeling to affect pathways governing mitochondrial turnover and intercellular exchange. Mitochondria are increasingly recognized as transferable organelles, exchanged between neurons and glial cells to support stress adaptation and metabolism^41^ . Importantly, mitochondrial fission, endolysosomal trafficking, and ER-mitochondria contacts play key roles in coordinating mitochondrial export and uptake^32–34,57,58^. We next asked whether spreading αS alters neuron–astrocyte mitochondrial transfer.

We examined mitochondrial exchange between iPSC-derived DNeurons and hiPSC-derived astrocytes (hAstrocytes) or human astrocytoma cells (CCF-STTG1, CCFs) using a non-contact co-culture system that permits intercellular transfer of organelle-derived vesicles without direct cell-cell contact (Fig. 4a). Mitochondria were differentially labeled in neurons and astrocytes, enabling quantitative assessment of bidirectional mitochondrial transfer under basal and stress-induced conditions. Neuronal mitochondria were labeled with MitoTracker Green FM and astrocytic mitochondria with MitoTracker Red CMXRos. Under basal conditions, neuronal mitochondria were detected within astrocytes after 14 h of co-culture, indicating spontaneous transfer (Fig. S4a). This was quantified by the neuronal mitochondrial fluorescence signal in astrocytes (Fig. S4b). Transfer efficiency further increased following oxidative stress induced by hydrogen peroxide (H_2_O_2_, Fig. 4b, c). Transferred neuronal mitochondria appeared as discrete puncta within recipient cells and localized to Lamp1-positive lysosomes (Fig. S4c), suggesting delivery for degradation. Conversely, astrocyte-derived mitochondria detected in neurons integrated into the neuronal mitochondrial network (Fig. S4d), supporting functional incorporation^41,64^.

**Figure 4.**
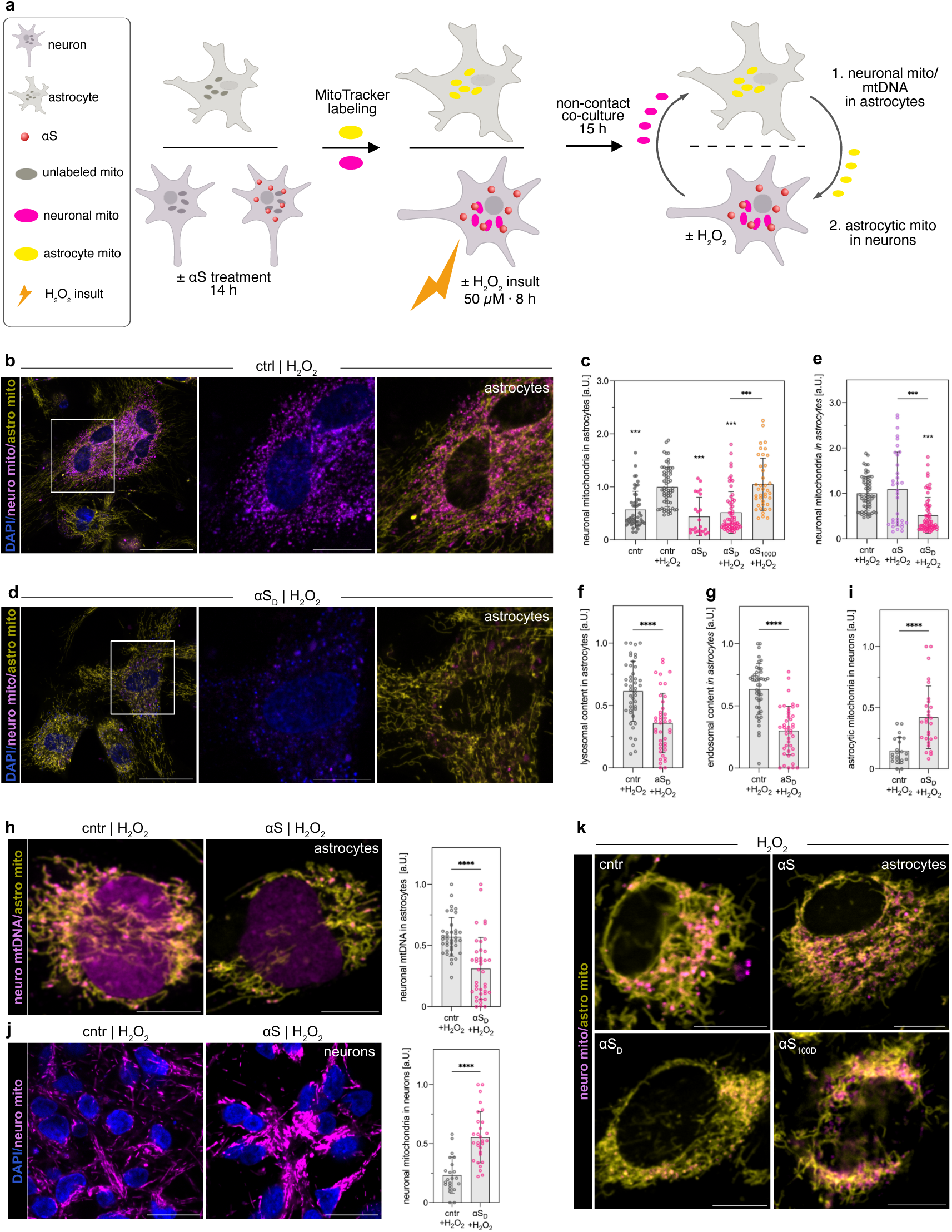
Spreading αS disrupts neuron-to-astrocyte mitochondrial transfer and alters intercellular mitochondrial exchange. a, Schematic of the non-contact neuron–astrocyte co-culture system used to assess the impact of αS on bidirectional mitochondrial transfer. Neuronal and astrocytic mitochondria were differentially labeled to quantify bidirectional mitochondrial exchange. Neuronal mitochondria (denoted neuro mito): magenta; astrocytic mitochondria (denoted astro mito): yellow; αS: red. b, Representative confocal images of astrocytes under H₂O₂ conditions showing uptake of neuronal mitochondria (neuro mito: magenta) into astrocytic cells (astro mito: yellow). Insets highlight transferred mitochondrial structures. c, Quantification of neuronal mitochondria detected in astrocytes under basal conditions and following oxidative stress (H₂O₂), in the presence or absence of αS_D_ or αS_100D_. αS suppresses both basal and stress-induced transfer of neuronal mitochondria to astrocytes, an effect that requires the αS C-terminus. LMM with random intercept (1|Replicate), Tukey HSD post-hoc on estimated marginal means, lmerTest/emmeans. *N=*2-4 biological replicates. d, Quantification of early endosomal intensity (Rab5) in astrocytes, following neuronal mitochondrial transfer to astrocytes under stress conditions. Astrocytic endosomal content is decreased in the αS_D_ condition. e, Quantification of lysosomal intensity (Lamp1) in astrocytes, following neuron-astrocyte mitochondrial transfer under stress conditions. Lysosomal content is decreased, indicating alterations in cargo marked for degradation. d, and e, Unpaired two-tailed Welch’s t-test; *N=*3 biological replicates. f, Representative images of astrocytes co-cultured with αS-treated neurons under H₂O₂ conditions, showing reduced neuronal mitochondria signal in astrocytes compared to control. g, Quantification of neuronal mitochondrial uptake by astrocytes in control and αS-treated conditions, demonstrating a significant reduction in transfer upon αS exposure in a concentration-dependent manner. Statistics as stated in c. h, Representative images and quantification of neuronal mtDNA (denoted neuro mtDNA: magenta) detected within astrocytes, confirming reduced transfer of neuronal mitochondrial material upon αS_D_ treatment. Unpaired two-tailed Mann-Whitney. *N=*2 biological replicates. i, Quantification of astrocyte-to-neuron mitochondrial transfer, showing increased transfer of astrocytic mitochondria into neurons upon αS exposure. Two-tailed Welch’s t-test; *N=*3 biological replicates. j, Representative images and quantification of neuronal mitochondria in control and αS-treated neurons upon stress. Neuronal mitochondrial signal is significantly higher for αS_D_, indicating retained neuronal mitochondria compared to control. Statistics as stated in i. k, Representative images comparing neuronal mitochondria signal in astrocytes following neuronal-to-astrocyte mitochondrial transfer. αS_100D_ does not reproduce the effect of αS_D_ on mitochondrial transfer. Data are presented as mean ± s.e.m.; each point represents an individual cell or field of view. * p < 0.05, ** p < 0.01, *** p < 0.001, **** p < 0.0001. Values normalized to within-replicate ctrl+H₂O₂ mean. Filled markers: no insult; open markers: +H₂O₂. Scale bars: c, f 50 µm; insets 20 µm, h, j, k 10 µm.

Strikingly, conditioning neurons with spreading αS markedly impaired this process. In the presence of αS localized to tri-organelle MAM-endosome sites, the transfer of neuronal mitochondria to hAstrocytes and CCFs were reduced by nearly twofold, even under H_2_O_2_ stress (Fig. 4c, d). The extent of this impairment scaled with αS concentration (Fig. 4e and Fig. S4e). This αS-dependent impairment of mitochondrial transfer is also evident under non-stress-induced conditions (Fig. S4b). Correspondingly, astrocytes co-cultured with αS-conditioned neurons exhibited a decrease in endo/lysosomal pool containing neuronal mitochondria (Fig. 4f, g), measured by immunostainings for Lamp1 and Rab5. This indicates reduced neuronal mitochondrial uptake and degradation by astrocytes. In parallel, quantification of neuronal mitochondrial DNA (mtDNA) using PicoGreen staining confirmed a significant reduction of neuronal mtDNA signal within astrocytes upon αS spreading (Fig. 4h). Interestingly, the reciprocal transfer of astrocytic mitochondria to neurons was enhanced (Fig. 4i). However, the neuronal mitochondria remained elevated compared to control where efficient transfer occurred (Fig. 4j). This bidirectional imbalance suggests that αS selectively interferes with the export of damaged neuronal mitochondria while maintaining or even promoting compensatory import of astrocytic mitochondria.

In a similar setup, αS_100D_ did not affect mitochondrial transfer from neurons to astrocytes (Fig. 4c, k, and Fig. S4f). These results indicate that the C-terminus of αS, and its associated disruption of calcium homeostasis^42,65^, is required for regulating mitochondrial export in neurons. Collectively, these findings reveal that spreading αS interferes with the normal directional exchange of mitochondria between neurons and astrocytes (Fig. 4c, k). We propose a twofold mechanism: (i) αS accumulation at MAM-endosome contact sites impairs mitochondrial fission and prevents damaged neuronal mitochondria from being transferred to astrocytes for degradation, and (ii) excess import of astrocytic mitochondria into neurons perturbs the stoichiometric balance of mitochondrial turnover and calcium buffering. Together, these effects may contribute to the progressive metabolic imbalance and neuronal vulnerability observed during the early stages of αS spreading and propagation in synucleinopathies.

## Discussion

The present study identifies organelle contact sites as an early and previously unrecognized target of spreading αS. Incoming monomeric αS preferentially localizes to mitochondria-ER-endolysosomal interfaces, where it remodels organelle communication and compromises MQC pathways. These findings suggest that recipient-cell responses to extracellular αS begin at discrete regulatory hubs that coordinate organelle dynamics and homeostasis, providing a mechanistic link between αS spreading and mitochondrial dysfunction in synucleinopathies.

A major implication of our findings is that extracellular αS does not distribute uniformly within recipient cells but instead accumulates at specialized contact-site populations. Organelle contact sites are increasingly recognized as signaling platforms^60^ that integrate endolysosomal sorting^66^, calcium homeostasis^31^, mitochondrial dynamics^32,34^, and stress responses^36,67^. The preferential recruitment of αS to tri-organelle interfaces therefore suggests that these structures represent vulnerable nodes through which relatively small amounts of transferred αS can exert disproportionate effects on cellular organization. Since αS used in this study is monomeric and below concentrations associated with aggregation^68^, our findings support a model in which contact-site dysfunction precedes the formation of classical αS pathology and may contribute to the earliest stages of disease progression.

Our data further suggest that the primary consequence of αS accumulation at contact sites is a loss of organelle plasticity. Dynamic remodeling of ER–mitochondria and endolysosomal contacts^61^ is essential for mitochondrial fission^32,34^, trafficking^58,66^, and quality control. We propose that αS stabilizes normally transient interactions, increasing organelle connectivity while reducing the flexibility required for adaptive remodeling. Such stabilization could impede the segregation and processing of damaged mitochondria, resulting in altered contact-site architecture and impaired MQC. Taken together, our findings suggest that mitochondrial dysfunction emerges not from an immediate loss of mitochondrial activity but from a progressive inability to remodel and eliminate damaged organelles.

The proteomic organization of αS-associated mitochondria supports this interpretation. Rather than exhibiting signatures of generalized mitochondrial failure, αS-associated mitochondrial fractions were enriched in pathways linked to organelle tethering, membrane remodeling, and metabolic regulation while displaying reduced representation of degradative trafficking and autophagic machinery. This pattern suggests a functional uncoupling between mitochondrial engagement and mitochondrial turnover. Mitochondria remain integrated within organelle communication networks yet become less efficiently connected to pathways responsible for their clearance. We therefore propose that spreading αS promotes a contact site-enriched but quality control-deficient mitochondrial state, characterized by persistent organelle connectivity and reduced degradative capacity. Importantly, these changes were restricted to αS-engaged mitochondrial populations, indicating that spreading αS selectively remodels discrete mitochondria. More broadly, this organization may reflect a cellular response to incoming αS. Although not directly tested here, such a model could contribute to the persistence of both dysfunctional mitochondria and intracellular αS within recipient cells.

The disruption of neuron-astrocyte mitochondrial transfer extends the consequences of contact-site remodeling beyond cell-autonomous dysfunction. Intercellular mitochondrial exchange is increasingly recognized as an important mechanism through which neural cells maintain tissue-level homeostasis during stress^36–41^. By reducing the transfer of neuronal mitochondria to astrocytes while enhancing the reciprocal delivery of astrocytic mitochondria, spreading αS appears to shift this system from mitochondrial disposal towards compensatory support. Such a response may temporarily preserve neuronal function but would be expected to favor the accumulation of damaged mitochondria within neurons. These findings suggest that defects in intercellular MQC may arise at early stages of αS spreading, before the emergence of neurodegeneration.

In summary, our findings support a model in which early αS spreading initiates mitochondrial dysfunction by altering the coordination of mitochondrial dynamics, quality control, and neuron–astrocyte communication. The spreading of αS shifts mitochondria toward a state of enhanced connectivity but impaired turnover. These events occur prior to aggregation and identify organelle contact sites as a potentially targetable vulnerability linking αS propagation to the earliest stages of synucleinopathies.

## Limitations and future directions

Several questions remain to be addressed. First, while our proteomic analysis identifies candidate pathways and proteins associated with αS-engaged mitochondria, the precise molecular interactions mediating αS recruitment to contact sites remain to be defined. Determining whether αS directly binds specific tethering complexes or alters membrane properties will be critical for understanding its mechanism of action. Second, the extent to which these findings translate *in vivo* and contribute to disease progression remains to be investigated. In particular, assessing how αS-mediated disruption of mitochondrial transfer impacts neuronal survival and circuit function. Third, our mitochondrial sorting strategy enables enrichment of αS-associated mitochondrial fractions, but it also has technical limitations. Contact-site proteins may be partially lost or redistributed during lysis, FAOS, and sample preparation for MS. Thus, the proteomic data should be interpreted as defining αS-associated mitochondrial populations rather than purified contact-site structures. Fourth, the mitochondrial transfer assay models vesicle-mediated exchange in a non-contact co-culture system. This design isolates transfer of organelle-derived material without direct cell–cell contact, but it does not capture tunneling nanotube–dependent mitochondrial transfer or other contact-dependent mechanisms. Therefore, future studies should test whether αS similarly disrupts mitochondrial exchange in contact co-culture systems and in animal models. Finally, the interplay between monomeric and aggregated αS species at contact sites remains unclear. The bidirectional imbalance in mitochondrial transfer suggests that targeting intercellular MQC pathways could be a therapeutic strategy. Enhancing mitochondrial export or restoring contact-site dynamics may help re-establish cellular homeostasis in the presence of spreading αS.

## Author contribution

M.B and E.F conceptualized the study and designed the experiments, E.F performed, analyzed and curated the data for most the experiments of the study, supervised by M.B and with contributions from: K.H analyzed and curated proteomic data, P.S.O performed the western blots, helped with stainings and supported neuronal differentiations, T.K prepared samples and performed mass spectrometry, and ran limma analyses, supervised by I.P, C.B supervised the FAOS, J. R performed imaging for stainings and FAOS preparation and differentiation of human astrocytes, L.B set up the pipeline for chromatic shift correction and RS-FISH spot-detection. M.B provided funding acquisition. E.F and M.B wrote the original draft and all authors participated in the review process.

## Supporting information

Hypotonic swelling on control untreated cells

Table of proteomic data with proteins of interest.

## Acknowledgements

We thank all members of the Birol laboratory for their support, insightful discussions, feedback, and continuous encouragement throughout this project. We thank Erich Wanker, Oliver Daumke, Johannes Broichhagen for valuable feedback, discussions, and advice during the development of this work. We also thank Johannes Broichhagen and Katja Simon for advice on the manuscript. We also thank the Siffrin laboratory for sharing astrocyte differentiation protocols. We acknowledge the support of the Systems Biology imaging platform at MDC, for microscopy assistance and technical expertise.

## Material and Methods

### Neuronal iPSC-derived cultures

The human iPSC line (BIHi005-A) was grown at 37 °C, 5% CO_2_ and 1% O_2_ in Essential 8^TM^ Flex Medium with supplements (E8+, ThermoFischer #A2858501) and passaged at 70%–80% confluency. For passaging, cells were treated with 0.5 mM EDTA (3 min), the solution was removed before the cells detached. Cells were detached and resuspended in E8+ and split 1:6 onto geltrex-coated wells of 6-well plates. Neural progenitor cells (NPCs) were generated from iPSCs by using the monolayer protocol as described in the STEMCELL’s Neural Induction kit (#A1647801; NIM+: 50 % Neural Induction media + 1x Neural Induction Supplement #A1647801; 50 % Advanced DMEM/F-12: #12634010 Thermofischer, 1 % Pen/Strep). For differentiation, iPSCs were split with Accutase (5 min), resuspended in DMEM/F-12, centrifuged (300x g, 3 min), resuspended in an appropriate volume of E8+ containing 5 μM rock inhibitor and seeded onto Geltrex-coated wells of 6-well plates with a plating density of 1x10^6^ cells. After 2-3 days, when cells reached 90 % confluency, differentiation to NPCs was initiated by performing daily medium change with NIM. After one week, cells were treated with Accutase (5 min), mixed with DMEM/F-12, passed through a 100 μm strainer, centrifuged (300× g, 3 min), resuspended in 3 mL NIM+ and seeded into wells of Geltrex-coated 6-well plates at a density of 1.5x10^6^ cells. NPCs were grown at 37 °C, 5% CO_2_ and 1% O_2_ and received fresh medium (NIM+) every day. Cells were passaged when 70%–80% confluence was reached, typically every 2–3 days, for 8-9 passages. For differentiation to midbrain dopaminergic neurons, NPCs were seeded onto PLO/Laminin coated wells of a 6-well plate at a density of 1.2x10^6^ cells and kept in NEM+ at 37 °C and 5% CO_2_ for one day post seeding. From day+1 after seeding, daily media change was performed for 6 days using the commercially available STEMDiff midbrain neuron differentiation kit (STEMCELL #10-0038). Following, midbrain dopaminergic neurons were prepared for maturation: Cells were washed with dPBS, split with Accutase, resuspended with DMEM/F-12 and centrifuged at (400x g, 5 min). Neurons were resuspended in a suitable volume of STEMdiff Midbrain Neuron Maturation Medium (STEMCELL #100-0041), seeded at a density of 7x10^4^ cells/cm^2^ on PLO/Laminin coated dishes and matured for 14 days. Half-media change was performed every 2-3 days, before subjecting to experiments. Neurons were kept in culture and used for a maximum of 14 days, with unchanged intervals of medium change. Neuronal cells were subjected to quality control (QC) immunofluorescence for neuronal and dopaminergic markers (as described below), and to QC calcium imaging to ensure functional maturation^69^.

### iPSC-derived human astrocyte differentiation

Human iPSC-derived astrocytes (h-astrocytes) were generated from NPCs as described by Alisch et al. with minor modifications^70,71^. Briefly, NPCs were generated from human iPSC line (BIHi005-A). Astrocyte differentiation was initiated after a minimum of 4 NPC passages, as successive NPC divisions progressively increase gliogenic competence. At the onset of differentiation, NPCs were detached with Accutase (5 min, 37 °C), resuspended in DMEM/F-12, passed through a 100 µm cell strainer, centrifuged (300 × g, 3 min), and plated at 5 × 10⁵ cells per well on 6-well plates pre-coated with Geltrex. The following day, culture medium was exchanged for astrocyte differentiation medium (ADM): DMEM base supplemented with 2 mM GlutaMAX-I (Thermo Fisher Scientific), 1% N-2 supplement (Thermo Fisher Scientific), 1% fetal bovine serum (FBS; Sigma-Aldrich), 20 ng/ml ciliary neurotrophic factor (CNTF; Miltenyi Biotec), and 1% penicillin/streptomycin. ADM was refreshed every 2–3 days. Upon reaching 80–90% confluency, cells were detached with Accutase and subcultured at a 1:2 to 1:3 ratio onto fresh Geltrex-coated surfaces. Cell number was recorded each passage to track proliferative activity. The progressive reduction in population growth served as an indicator of post-mitotic astrocyte maturation. Differentiation was continued for up to 6 weeks or until proliferation arrested, at which point cells were seeded at high density without further dilution. Forth on, h-astrocytes were split 1x per week without further dilution but compensating for any loss of cell count. Astrocyte identity was verified by immunofluorescence confirming expression of GFAP and S100β.

### Co-culture assays to monitor mitochondria transfer

24-well plates were decorated with three paraffin wax pillars per well that later served as physical spacer for co-culture cell-growth surfaces. Neurons were seeded on PLO/laminin-coated wells of these 24-well plates, astrocytes on Geltrex-coated ⌀12 mm coverslips. αS-treated (14 h) and untreated midbrain dopaminergic neurons were incubated with 100 nM MitoTracker Green FM (Thermofisher, #M7514) or 1:1000 dilution of PicoGreen (30 mins), immortalized and hiPSC-derived astrocytes were labeled with 50 nM MitoTracker Red CMXRos (Thermofisher, #M7512, 30 mins, 37 °C). After protein treatment and mitochondria labeling, cells were washed with warm dPBS. Neuronal insult was induced by treating neurons with 50 µM H_2_O_2_ for 8h post MitoTracker labeling but prior to co-culture assembly. For co-culturing, astrocyte-holding glass coverslips were placed facing down onto paraffin pillars of neuron-holding wells, creating a non-contact co-culture. Co-cultures were topped up with fresh neuronal maturation medium reaching a total of 900 µl without removing the pre-existing medium of the neuronal portion of the co-cultures. To measure mitochondria transfer, co-cultures were incubated for 15 h (37 °C, 5% CO_2_) before imaging. For imaging, co-culture medium was replaced with BrainPhys Imaging Optimized Medium (STEMCELL #05796) and cells were left to recover (10 min). When applicable, astrocytic portion of co-cultures was fixed and stained with LAMP1 and Rab5 to assess for changes in the endolysosomal pool in astrocytes. Imaging was performed on a Dragonfly spinning disk confocal microscope (Andor, Oxford Instruments) or a PicoQuant MicroTime 200 time-resolved fluorescence system as described below. Mitochondrial transfer was quantified in two co-culture experimental designs: 1. In co-cultures with MitoTracker-labelled neurons and astrocytes, mitochondrial uptake was measured in both directions (neuronal mitochondria uptake by astrocytes, astrocytic mitochondria uptake by neurons), 2. In co-cultures with PicoGreen-labelled neurons and MitoTracker-labelled astrocytes, uptake of neuronal mtDNA in astrocytes was quantified.

### Cell culture

The following human cell lines were used: SH-SY5Y (RRID: CVCL_0019) and CCF-STTG1 (RRID: CVCL_1118). SH-SY5Y and CCF-STTG1 were obtained from ATCC (ATCC-CRL-2266 and ATCC-CRL-1718). The experiments were performed on the original cell lines. SH-SY5Y cells were cultured in Dulbecco’s Modified Eagle’s Medium (DMEM) plus 10% fetal bovine serum (FBS), 50 U/ml penicillin, and 50 μg/ml streptomycin. CCF-STTG1 cells were grown in RPMI with the same supplementation. Cells were grown at 37°C under a humidified atmosphere of 5% CO_2_. Cells were passaged upon reaching ∼ 95% confluence (0.05% Trypsin-EDTA, Life Technologies, Carlsbad, CA), propagated, and/or used in experiments. Cells used in experiments were pelleted and resuspended in fresh media lacking Trypsin-EDTA before replating.

For imaging of SH-SY5Y cells, cells were grown in eight-well Ibidi chambers (μ-Slide 8 Well high, #1.5 polymer, ibiTreat, Ibidi GmbH, Germany, #80806). Chambers were seeded with 4x10^4^ cells/well and cultured for 48 h before beginning experiments. For cellular uptake experiments, ATTO700-/Alexa647-labeled αS was added to fresh cell media to a final concentration of 80nM (for αS) and 250nM (for αS_D_ or αS_100D_). Following incubation (0 to 24 h, indicated for each experiment), the media was exchanged to remove non-internalized αS and cells were allowed to recover (10 min) before acquiring images.

### αS purification

αS was expressed in E. coli BL21 cells; for αS, BL21 stocks containing the N-terminal acetyltransferase B (NatB) plasmid with orthogonal antibiotic resistance were used. αS constructs were expressed in 0.5 l growths in LB media at OD600 ≈ 0.4-0.5 by induction with 320 µM IPTG for 2 h at 37°C. Cells were pelleted and resuspended in fresh lysis buffer (25 ml per 0.5 l; 40 mM NaOH, 20 mM Tris pH 8.0, 1 mM EDTA, 1mM PMSF, 0.1% Triton X-100), supplemented with 1 mM PMSF and a cOmplete protease inhibitor cocktail tablet. The lysate was then flash-frozen in liquid nitrogen and stored at -80°C until use. The purification of αS full length protein and αS_1-100_ was carried out as previously described^72,73^, with minor modifications. Briefly, two ammonium sulfate cuts were used (0.116 g/mL and 0.244 g/mL) with αS precipitating in the second step. The pellet was resolubilized in Buffer A (25 mM Tris [pH 8.0], 20 mM NaCl, 1 mM EDTA) with 1 mM PMSF and dialyzed against Buffer A to remove ammonium sulfate overnight. Dialyzed samples were filtered through a 0.22 μm filter, loaded to an anion exchange column (GE HiTrap Q HP, 5 ml) and eluted with a gradient to 1 M NaCl. αS elutes at approximately 300 mM NaCl. Fractions containing αS were pooled, concentrated using Amicon Ultra concentrators (3,000 Da or 10,000 Da MWCO) and buffer exchanged to Buffer C (25 mM Tris [pH 8.0], 100 mM NaCl, 1 mM EDTA and 0.5 mM TCEP). Concentrated samples were then loaded to a size exclusion column (GE HiLoad 16/600 Superdex75) and eluted at 0.5 ml/minute in Buffer C containing 1 mM TCEP overnight. The next day, fractions containing αS were confirmed by running an SDS-gel, pooled and concentrated to then proceed with the labeling on the same day. Unlabeled αS fractions were pooled, concentrated, snap-frozen at a concentration of ∼30 μM and stored at -80 °C.

### αS labeling

For site-specific labeling of αS, a cysteine was introduced at residue 9 (S9C point mutation) For labeling reactions, freshly purified αS (typically 300-500 μl of ∼100 μM protein) was incubated with 1 mM dithiothreitol (DTT) for 30 min at RT to reduce cysteines. The protein solution was buffer exchanged into Buffer D (20 mM Tris (pH 7.4), 50 mM NaCl) with additional 6 M guanidine hydrochloride (GdmCl) and concentrated to 1 mL using Amicon Ultra concentrators (3,000 Da or 10,000 Da MWCO). The protein was incubated with 4-fold molar excess ATTO 700 or Alexa647 maleimide (Invitrogen) overnight at 4°C under stirring conditions. The labeled protein was buffer exchanged into fresh Buffer D and concentrated using Amicon Ultra concentrators (3,000 Da or 10,000 Da MWCO). Access dye and GdmCl was removed by passing the solution through two coupled HiTrap Desalting Columns (GE Life Sciences, Pittsburgh, PA) equilibrated with Buffer D. Fractions containing labeled αS were pooled, diluted to ∼30 μM with Buffer D, snap-frozen and stored at -80 °C. Concentration and momomeric state of labelled protein was verified using fluorescence correlation spectroscopy (FCS) on a PicoQuant MicroTime 200 time-resolved fluorescence system after freezing and approximately once very calendar quarter to check for degradation or aggregation. Alexa647-labeled αS was utilized for FACS and MS/proteomic studies, all other experiments were performed with ATTO 700-labeled αS.

### Transfection and nucleofection of SH-SY5Y cells

SH-SY5Y cells were seeded onto 8-well ibidi chambers and cultured to ∼70% confluency prior to transfection. Cells were transfected with SPLICS Mt-ER short P2A (addgene: Plasmid #164108) using Lipofectamine 3000 (Invitrogen #L3000001) according to the manufacturer’s instructions, with 250 ng plasmid DNA per well. t 30-36 h post-transfection, culture media was exchanged with fresh media prior to αS-treatment.

Stable overexpression cell lines were generated by nucleofections of SH-SY5Y cells using the Amaxa SF Cell Line 4D-Nucleofector X Kit L (Lonza #V4XC-2024) and the Lonza 4D-Nucleofector System (Core Unit AAF-1003B, X Unit AAF-1003X), by following the Amaxa 4D-Nucleofector Protocol for SH-SY5Y using a final concentration of 250 ng of plasmid. αS-SNAP plasmid was generated (pCMV-αS-SNAP, SNAP-tag cloned C-terminally in pCMV- αS backbone), mCherry-VAPB was purchased from addgene (addgene: Plasmid #108126). Cells were selected by treatment with 300 µg/ml geneticin (G418-Sulfat), expression was verified with fluorescence imaging.

### Hypotonic Cell Treatment

To generate large intracellular vesicles (LICVs) from mitochondria, SH-SY5Y cells conditioned with and without αS were treated with hypotonic media (10% DMEM in water, pH ∼ 7), incubated at 37 °C with 5% CO2 for 10 min to allow for Mito-LICV generation, and then imaged as previously shown^61^. Neurons were treated with hypotonic LICV media (50% DMEM in water, pH ∼ 7), incubated at 37 °C with 5% CO2 for 10 min to allow for mito-LICV generation, before image acquisition. Following hypotonic treatment, SH-SY5Y cells remained stable in the imaging chamber for approximately 60 min, whereas neurons remain viable for less than 30 min. Following longer exposure to hypotonic solution, cells disintegrate, burst and/or detach from the dish. Pre-maturely detached or lysed cells were not imaged or utilized in this study.

### Live cell fluorescent organelle dyes and stains

Cells were treated with 20 nM MitoTracker Red CMXRos (Thermofisher, #M7512) for 30 mins before imaging. For mitochondrial transfer assays, neurons were treated with 100 nM MitoTracker Green FM (Thermofisher, #M7514), astrocytes with 50 nM MitoTracker Red CMXRos for 30 min prior to H_2_O_2_ treatment and co-culturing. mtDNA was labelled in neurons using PicoGreen dsDNA reagent (Quant-iT™, P11496, Thermo Fisher Scientific) at a 1:1000 dilution from the concentrated stock solution. For colocalization of αS and endolysosomes, cells were treated with 400 nM LysoSensor Green DND-189 (Thermofisher, #L7535) for 30 mins before imaging. Prior to live imaging, cells imaged on the ANDOR Dragonfly system were stained with 2 µg/ml Hoechst for 5 min at 37 °C before exchanging media for fresh imaging media.

### Immunofluorescence

For immunofluorescence, cells were fixed for 15 min with 4% PFA (w/v) (SH-SY5Y) or 4% PFA and 4% sucrose (dopaminergic neurons and astrocytes), supplemented with 0.1% glutaraldehyde. Cells were washed 3x with dPBS for 5 min after fixation and every subsequent step. Cells were permeabilized with 0.2% Triton X-100 in dPBS for 5 min, then blocked for 1 h at room temperature with 0.5% BSA and 10% normal goat/donkey serum. Primary staining was performed at 4 °C in 3% BSA overnight, shaking. Primary antibodies used were as follows: for endolysosomal pool characterization Rab5 (1:200; monoclonal rabbit anti-Rab5 [ERP21801], abcam #ab218624) and LAMP1 (1:200, monoclonal mouse anti-LAMP1 [H4A3], Developmental Studies Hybridoma Bank (DSHB) #AB_2296838); for QC of midbrain dopaminergic neuronal, Tyrosine Hydroxylase (1:200; polyclonal rabbit anti-TH, Abcam #ab152), FOXA2 [HNF-3β] (1:200; polyclonal goat anti-TH, R&D Systems #AF2400), TUJ1 (1:500; monoclonal mouse anti-Tubulin-β3, Biolegend #801201); for QC of iPSC derived h-astrocytes, GFAP (1:200; polyclonal rabbit anti-GFAP, Agilent #Z033429-2), S100-beta [EP1576Y] (1:100; monoclonal rabbit anti-S100 beta, abcam #ab52642). After washing samples 3x 10 min in dPBS, secondary antibody staining was performed in 0.5% BSA solution for 1-2 h at room temperature (1:800-1000; anti-rabbit Alexa647 abcam #ab150075; anti-mouse Alexa488 Life Technologies #A21202; anti-rabbit Alexa594 Life Technologies #A11012; anti-goat Alexa680 Invitrogen #A21084). Samples were washed 3x 10 min in dPBS, during the last wash DAPI was added at 0.4 µg/ml.

### Cell imaging

Live cell imaging was performed at 37 °C and 5% CO_2_. For hypotonic cell swelling assays in dopaminergic neurons, and for quantitative analysis of mitochondrial transfer, live cell samples were imaged on an ANDOR Dragonfly spinning disk confocal microscope (Andor, Oxford Instruments, Abingdon, UK) equipped with a 63 × Plan-Apo/1.1-NA glycerol-immersion objective. Across conditions within the same experimental replicate, excitation/emission (ex./em.) and gain settings for the channels were kept constant: DAPI, Hoechst - 405/450; MitoTracker Green FM, SPLICS – 488/525; mCherry, MitoTracker Red CMXRos – 561/600; ATTO700-αS, Alexa647-αS – 637/700.

All other imaging was carried out on a PicoQuant time-resolved fluorescence system with a TCSPC MicroTime 200 module, based on an inverted Olympus IX73 microscope (Olympus, Tokyo, Japan) with a 60X Plan-Apo/1.4-NA water-immersion objective and 479/560/680 nm excitation lasers. Time-gated pulsed interleaved excitation (PIE at 26.76 MHz) was conducted throughout all acquisitions, with a triple-line beamsplitter directing signals to SPAD detectors. Images were acquired at a frame size of 428 × 428 pixels and a dwell time of 150 µs, resulting in a frame-acquisition speed of 55 s. For time-lapse imaging, these settings were maintained continuously for 30 min, interrupted only by re-adjusting the z-plane if necessary. Images acquired with this instrument were in fluorescence lifetime mode but were integrated in the SymPhoTime 64 software (PicoQuant, Berlin, Germany) to obtain intensity-based images, auto-exported as TIFF files. Image processing and analysis were performed either in Fiji^74^, utilizing Fiji in-built plugins, custom made Fiji-based macros, or python-based scripts.

### Western blots

For protein extraction, cultured cells were scrapped off and lysed with RIPA buffer (25mM Tris-HCl pH 7.6, 150 mM NaCl, 1% NP-40, 1% sodium deoxycholate, 0.1% SDS) with 1x Protease Inhibitor Cocktail (Sigma Aldrich) for 45 min on ice, vortexing every 5 min and sonicating 3x for 1 min in a water bath sonicator. To remove cell debris the solution was centrifuged for 20 min at 10,000 rpm at 4 °C. Protein concentration was calculated using the BCA Protein Assay Kit (Thermofisher). Protein samples (40 and 20 μg) were mixed with 4x Laemmli Buffer and boiled at 95°C for 5 min. Samples were loaded on 8 – 16 % Tris-Glycine Plus WedgeWell precasted gels (Invitrogen), separated by SDS-PAGE and transferred to 0.45 μm PVDF membranes (Sigma Aldrich). Membranes were blocked for 2 h at room temperature with 5% BSA in 1x TBST (TBS, 0.1 % Tween20) and incubated overnight at 4 °C with primary antibodies diluted in the same solution against αS (rabbit anti-alpha + beta Syn, 1:1000, ab51252, Abcam) and vinculin (mouse anti-vinculin, 1:1000, V9131, Sigma Aldrich). Membranes were washed, incubated for 1 h with the appropriate secondary antibody conjugated to horseradish peroxidase (Cell Signaling) at room temperature, and the immunoblots were developed using ECL Western Blotting Detection Reagent (Cytiva). The Gelanalyzer Software was used for imaging and analysis.

### Sample preparation for FACS sorting

14 h prior to experiment, cells were treated with Alexa647 labelled αS or αS_D_. Cells were harvested, pelleted and samples kept on ice. Cells were lysed with custom-made mild lysis buffer (10 mM Tris-HCl, 10 mM NaCl, 3 mM MgCl₂, 0.01% NP-40, 1 mM DDT, 1x protein inhibitor cocktail +EDTA, 1 µM PMSF, in nuclease-free H_2_O), and kept on ice for 10 min, resuspending every 2 min. Samples were sonicated (one cycle, 5 s) in a sonicator bath (Bioruptor Next Gen, Diagenode #B01020001) and filtered through 30µm strainer, rinsing strainer once with washing buffer (dPBS, 1x protein inhibitor cocktail + EDTA, 1µm PMSF) to collect residues. Whole cell lysates were incubated on ice for 5min prior to staining with 100 nM MitoTracker Green FM (Invitrogen M7514) for 10 min on ice. Lysates were then filtered through a 20 µm strainer into Polypropylen FACS tubes and subjected to FACS.

### Fluorescence-Activated Organelle Sorting (FAOS)

Whole-cell lysates were sorted on a BD FACSDiscover S8 cell sorter (BD Biosciences) equipped with BD CellView Image Technology and BD SpectralFX Technology, operated via FACSChorus software (v6.2). From the standard five-laser configuration and three imaging detection channels, MitoTracker Green FM was excited by the 488 nm laser, detected spectrally and imaged on the CellView imaging detector, αS_A647_ was excited by the 637 nm laser and detected. Spectral unmixing was performed using the FACSChorus unmixing algorithm with single-stained or unstained controls (unstained control, unstained αS_A647_, unstained samples treated with unlabeled (dark) αS, dark αS stained with MitoTracker Green). Sorting was performed with the 85 µm nozzle configuration (35 psi, 57 kHz) in Purity mode at a FACSChorus flow rate setting of 10–20, adjusted according to sample concentration. Sample and collection tubes were maintained at 4 °C and 400 rpm. Gating strategy for FAOS of αS-associated mitochondria. A four-gate hierarchy (P1–P4) was applied to isolate individual mitochondrial entities on the basis of MitoTracker Green fluorescence, axial light loss, and BD CellView imaging features (eccentricity, radial moment, and centre of mass), followed by separation into αS_A647_-positive (αS**^+^**) and -negative (αS**^-^**) mitochondrial populations. Sequential gates were applied to whole-cell lysates processed for FAOS on the BD FACSDiscover S8. Mitochondrial events (P1) were identified by MitoTracker Green FM fluorescence in combination with BD CellView imaging, selecting morphologically small, circular structures by axial light loss and imaging parameters. Singlet events were isolated from P1 (P2) by exclusion of elongated and aggregated events based on eccentricity and radial moment. Centered events (P3) were selected from P2 by center-of-mass gating to retain only events fully contained within the imaging field of view. From P3, αS-associated (αS⁺) and αS-negative (αS⁻) mitochondrial populations were resolved spectrally by A647 fluorescence (P4).

Sorting was continued until the entire lysate volume was processed rather than to a fixed event target, to maximize recovery of αS-associated mitochondrial events. Across samples, 107,000–190,000 αS**^−^**mitochondrial events and 2,500–5,500 αS**^+^** events were collected from αS-treated samples, and 100,000–120,000 total mitochondrial events from control samples. Variation in total event counts reflected differences in sample loss during filtration and lysis efficiency. Samples were collected simultaneously into 1.5 mL LowBind Ependorf tubes pre-filled with washing buffer, maintained at 4 °C during sorting and flash-frozen immediately after. Data were exported in FCS 3.2 format and analyzed in FlowJo (v.10) with the BD CellView Lens Plugin for concurrent analysis of imaging and spectral parameters.

### Sample processing and Mass Spectrometry

For each sample with 3000 mitochondrial events or less, the whole sample was taken for subsequent processing. Otherwise, a fraction of the sample, equivalent to 3000 mitochondrial events, was used to ensure that all subsequent steps were performed on the same amount of starting material. The samples were spun down (18,000 x g) for 15 min at 4°C and the supernatant removed. 15 µl of 0.1% N-Dodecyl-β-D-maltoside was added in the presence of 100 mM triethylammonium bicarbonate buffer, 5 mM tris(2-carboxyethyl)phosphine) and 20 mM chloroacetamide. Mitochondria were lysed by sonication (10 min, 30 sec on 30 sec off at 4 °C) using a Bioruptor (Diagenode, Seraing, Belgium). Samples were incubated overnight with a mixture of Lys-C (20 ng) and trypsin (20 ng) at 37 °C. The digestion was quenched with formic acid and then cleaned using Evotip Pure C18 tips (EV2013, Evosep, Odense, Denmark). Tips were eluted with 35% ACN in 0.1% formic acid. Samples were dried and resuspended in 5.4 µl of loading buffer. 5 µl of the sample was injected.

Peptides were analyzed using a label-free data-independent acquisition (DIA) workflow on an Orbitrap Astral Mass Spectrometer (Thermo Fisher Scientific, Waltham, MA USA) coupled to a Vanquish Neo UHPLC system (Thermo Fisher Scientific) operating in nano-flow mode. Peptides were separated on a 75 μm x 20 cm analytical column (packed in-house with 1.9 μm C18 resin; Reprosil-Pur 120 C18-AQ, Dr. Maisch) applying a flow of 250 nl/min over 17.5 min using solvent A (0.1% FA, 3% ACN) and solvent B (0.1% FA in 90% ACN). Briefly from 0 to 1 min solvent B was increased from 2 to 7%, followed by an increase to 20% and 30% solvent B, in 10 and 7.5 min, respectively. Solvent B was ramped up to 60% B over 2 min, followed by column washing at 90%B. Total run time was 25 min. The column was then equilibrated at high flow with a column equilibration factor of 3. For the DIA method, MS1 acquisitions were performed in the Orbitrap analyzer with a scan range from 380–1100 *m*/*z* at a resolution of 240,000 a normalized Automatic Gain Control (AGC) of 500% and a maximum injection time of 5 ms. MS2 acquisitions were performed using a range from 400-800 *m*/*z*, an isolation width of 4 Th, a maximum injection time of 8 ms, a normalized AGC of 500% at 25% collision energy.

### Proteomic data analysis and representation

Raw data were processed in Spectronaut 20^75^ (Biognosys, build 20.2.250922.92449) against the reviewed *Homo sapiens* UniProt database (2023-05; 20,407 entries) using default settings, with cross-run normalization enabled and imputation disabled. Trypsin and endopeptidase Lys-C were specified as digestion enzymes. All downstream analyses were performed in R (v. 4.5.2; https://www.r-project.org/). For each comparison, proteins were retained if quantified in the relevant sample set (pairwise: αS^+^ vs. control, αS^-^ vs. control; pooled: αS^+^/αS^-^ vs. control). Sample clustering was assessed by principal component analysis on log2-transformed protein intensities. Differential abundance between αS^+^, αS^-^ and control was assessed using moderated t-tests (pairwise) in limma^76^ (v. 3.66.0) , with empirical Bayes moderation using an intensity-dependent mean–variance trend. P-values were adjusted by the Benjamini–Hochberg procedure; proteins were considered differentially abundant at FDR < 0.05 and |log2 fold-change| > 1. Gene symbols were mapped to Entrez IDs using biomaRt^77^ (v. 2.66.0). Gene set enrichment analysis (GSEA) was performed on individual and pooled comparisons using clusterProfiler^78^ (v. 4.18.1) against Gene Ontology, Reactome and KEGG term sets. Differentially expressed proteins (DEP) were filtered by eulerr package (v. 7.0.4; https://doi.org/10.32614/CRAN.package.eulerr) for shared, as well as unique DEPs within αS^+^ vs. cntr and αS^+^ vs. αS^-^ comparisons. And then both shared and unique DEPs were subjected to over-representation analysis (ORA).

Terms appearing in both comparisons’ unique lists with non-overlapping gene fingerprints were excluded. For both GSEA and ORA, enrichment outputs were filtered to a custom curated list of existing mitochondria- and organelle-related terms in databases across GO, Reactome, KEGG and MitoCarta^79^. Redundancy of Gene Ontology (GO), Reactome Pathway (RP) and KEGG terms was reduced by semantic similarity and hierarchical relationships, and representative parent terms were selected to summarize related biological child-terms. For Cnet and heatmaps, pairwise Jaccard similarity of leading-edge protein sets (threshold 0.60) was performed. Retained terms were filtered manually and visualized as dot plots, or as cnet plots and heatmaps plots using ggplot2 (v. 4.0.0), igraph (v. 2.2.1) and pheatmap (v. 1.0.13).

In parallel, a curated list of 207 proteins of interest (177 detected in this dataset) across 13 functional categories (Supplementary Table S1) was compiled from literature, spanning mitochondrial morphology and dynamics, trafficking, quality control, lipid metabolism and transport, organelle contact sites, and intra-and intercellular cargo trafficking. Differential abundance results were visualized as volcano plots using ggplot2, with proteins of interest color-coded by manually defined functional category over a background of all detected proteins. Protein names were annotated only for proteins explicitly discussed in the main text; all remaining proteins of interest are shown as colored points without labels.

### Data analysis

#### Chromatic aberration correction

To correct for chromatic aberration between fluorescence detection channels on the PicoQuant System, image registration was performed using a custom Python-based pipeline. Registration was performed using the SimpleITK library^80^ (version 2.4.1), employing a translation transform optimized by the Mattes Mutual Information metric computed over 500 histogram bins, thereby enabling calibration-free image registration. This intensity-based metric quantifies the statistical dependence between image channels, allowing registration parameters to be derived directly from the image data without an external fiducial calibration target, such as fluorescent beads. Optimization was performed using a Regular Step Gradient Descent optimizer with a learning rate of 1.0, a minimum step size of 1×10⁻⁵, a maximum of 1,000 iterations, and a gradient magnitude tolerance of 1×10⁻⁸. During optimization, B-spline interpolation was applied to the moving image; the final registered channel was resampled onto the fixed image grid using linear interpolation.

For each three-channel image, channel 1 served as the fixed reference, to which channels 2 and 3 were independently registered. Pixel-size information lost during registration was recovered from the original images. Registered multichannel images were split into single-channel TIFF files for downstream analysis, with spatial calibration metadata preserved throughout. Chromatic aberration correction spatially transforms pixel data via interpolation; therefore, distance-based colocalization and other analyses requiring sub-pixel spatial precision were excluded. Colocalization analysis was therefore restricted to object-based approaches.

#### Fluorescence intensity line profiles

Fluorescence line intensity profiles (Fig. 1 and Fig. S1), line segments (6-10 µm) were drawn across ROIs, and intensity values for each channel were extracted using the Fiji in-built plot profile function. Profiles from multiple regions (n = 2-3) were stacked and plotted using a custom Python script. Channel intensities were normalized using scaling factors to visualize spatial relationships, since absolute intensity values were not compared between channels.

#### Segmentations

Cell-area segmentations were either performed using Cellpose^81,82^ (v3.1.1.) or in Fiji. Neuronal somata were segmented using the built-in cyto3 model with automatic diameter estimation, and predicted masks were manually curated in the Cellpose GUI to correct segmentation errors. Cellpose mask files (.seg.npy) were converted to Fiji-compatible ROI archives (.zip). For each image, the area of the cell soma was measured by combining all individual soma ROIs into a compound selection in Fiji and recording the total area. In Fiji, saturation-threshold images were converted to binary masks, and Fiji’s built-in binary operations were applied for refinement using the same combination across all images from the same experiment. Masks of cell somata were used to calculate the area of individual cells or the total cell area per image. Per-cell area values were used to normalize MAM-density (Fig. 1j), total cell-area per image was used to normalize average intensities of LAMP1 and Rab5 measurements (Fig. 1f and Fig. 4d, e), assess mitochondria morphology (Fig. 2), and for quantifications of mitochondrial transfer assays (Fig. 4 and Fig. S4). Mitochondria were segmented manually in Cellpose by tracing each mitochondrial object in the Cellpose GUI annotation interface; automated prediction was not used to delineate partially or fully fragmented mitochondrial networks accurately.

#### MAM properties and αS-MAM localization

MAM contact sites and αS puncta were detected using the radial symmetry-based spot detection algorithm RS-FISH^83^ (Fiji Plugin). To obtain puncta areas for size quantification, detected coordinates were expanded into circular ROIs using a custom-built Fiji macro that performs full-width-at-half-maximum (FWHM)-based radial profiling with adaptive thresholding. Size limits of 0.15–0.70 µm radius were applied for MAM puncta and 0.20-80 µm based on manually measured size ranges. Puncta exhibiting anisotropic expansion profiles (max./min. radius ratio >2.5) were flagged and excluded from size analysis. Measurements displaying artifactual clustering at size-limit cutoffs (9.8% of data) were removed prior to statistical analysis and plotting.

MAM density was calculated as the number of RS-FISH-detected puncta per cell area, expressed as puncta per 1000 µm². MAM size (radius [µm] and area [µm^2^]) was quantified from FWHM-expanded ROIs. Statistical comparisons were performed using linear mixed models (LMM) with condition as a fixed effect and biological replicate as a random effect, implemented in R (lmerTest package). Effect sizes (Cohen’s d) and estimated marginal means with 95% confidence intervals were computed; skewness (γ) was reported per condition.

Colocalization between αS and MAM puncta was quantified using object-based analysis on pre-segmented ROIs generated by the FWHM-based approach. Area-overlap coefficients (M1, M2) and Jaccard index were calculated. Statistical significance was assessed by Monte Carlo randomization (1000 iterations). Observed values were compared to randomized distributions using Student’s t-test. Comparisons between conditions were performed using one-way ANOVA (M1) or Kruskal-Wallis test (M2), and figures were generated in GraphPad Prism (v11.0.0).

#### Endolysosomal pool

For endolysosomal pool quantifications in fixed cells (Fig. 1f in SH-SY5Y cells, Fig. 4d, e in CCF-STG1), total intensities of LAMP1 and Rab5 antibody signal were measured (Fiji, RawIntDen) only taking into account ROIs within cell-somata masked areas. For each image and channel, total fluorescence intensities were plotted per total cell area, separately or as ratio. For live imaging data, individual endolysosomal ROIs were manually generated in SymphoTime from the lifetime-based images of LysoSensor channel, exported as bitmaps and converted to binary masks from the RGB photon count image in Fiji. For αS-treated conditions, endolysosomal puncta were classified as αS-associated (spatially overlapping with αS signal) or αS-distal (remaining puncta) and treated individually or pooled. For untreated control cells, all endolysosomal puncta were pooled as a single population. Mean fluorescence intensity per ROI was extracted in Fiji as the integrated density divided by ROI area (Fiji “Mean” value), providing a size-independent measure of LysoSensor signal per organelle.

#### Mitochondrial morphology analysis

Mitochondrial morphology and network connectivity was quantified from single-plane using a pipeline combining manual segmentation in Cellpose with automated morphometric analysis in Fiji (ImageJ v2.14.0). Images from SH-SY5Y, αS-SNAP cell lines, and neurons were processed with cell-type-specific parameters, held constant across all conditions and experiments of a given cell line. **Pre-processing.** Images were background corrected: For SH-SY5Y and αS-SNAP images, a rolling-ball background subtraction was applied in Fiji (Subtract Background, radius = 15–30 px, optimised per experiment and held constant across all conditions within the experiment). For neuronal images, background was removed by subtraction of a Gaussian-blurred copy of the image. Cell-soma and mitochondrial masks were generated in Cellpose as previously stated.

#### MitoAnalyzer pipeline

Mitochondrial ROIs were refined by clearing pixel intensities outside the ROI boundaries in Fiji, producing per-image ROI-restricted intensity images. These were processed with the Mitochondria Analyzer plugin for Fiji^84^ (v2.3.0) using the 2D Threshold and 2D Analysis modules in batch mode via an in-house Fiji macro. Thresholded images were hole-filled (Fiji’s Fill Holes (Binary/Gray)), after which 2D Analysis was run to extract per-image morphometric descriptors. Two metrics were used for downstream analysis: (i) the mean mitochondrial branch length per image, calculated as the total branch length divided by the number of branches, (ii) the mean mitochondrial aspect ratio per image, obtained from ellipse-fitted major/minor axis ratios across all detected mitochondrial objects and used as a measure of elongation. **Analyze Particles pipeline.** To quantify mitochondria vesicle density per cell area, a complementary particle-based analysis was performed on binary mitochondrial masks generated from the Cellpose segmentations. For SH-SY5Y and αS-SNAP cultures, binary masks were additionally filtered with three iterations of the Fiji Despeckle (3 × 3 median) filter to remove isolated-pixel noise. This step was not applied to neuronal samples, as median filtering led to erosion of their finer tubular structures. Particle analysis was performed using Fiji’s Analyze Particles function restricted to the combined cell-soma ROI, with a circularity range of 0.50–1.00 and a size range of typically 0.3–∞ µm² (reference: smallest to largest vesicle in control sample, varied between data sets). The per-image particle count was then divided by the corresponding per-image cell-soma area to yield the mitochondrial vesicle density (vesicles per µm²). **Data handling and pooling.** All three metrics were computed on a per-image basis, with one image contributing one data point per metric per biological replicate. For mitochondrial vesicle density and mean branch length, data were min–max normalised across all conditions within each biological replicate prior to pooling. Aspect ratio remained unnormalized. **Timelapses.** Timelapses were generated using the concatenate function in Fiji and time stamps added with Fiji’s Time Stamp function.

#### Mitochondrial transfer assay

For each image, total fluorescence intensity of the transferred-label channel (MitoTracker Green for astrocytic uptake of neuronal mitochondria, MitoTracker Red CMXRos for neuronal uptake of astrocytic mitochondria, or PicoGreen for astrocytic uptake of neuronal mtDNA) was measured in Fiji within cell-soma masks (generated as described above) and normalized to the total nuclei count per image, yielding a single intensity-per-cell value per image. Outliers were detected and removed using IQR fences and median absolute deviation (MAD) as described. Within each biological replicate, values were expressed relative to the mean of the H₂O₂-treated reference condition (in GraphPad Prism: descriptive statistics and transform function). Normalized values were then pooled across replicates for visualization and statistical comparison. The MitoTracker Green/Red co-culture experiments were performed in *n* =3–5 biological replicates, and the PicoGreen/MitoTracker Red co-culture experiments in *n*=2 biological replicates.

### Image upscaling

Higher magnification representative images of αS-mitochondria-endolysosome and αS-mitochondria-SPLICS localization were upscaled to 500x500 pixels, representative images αS-SNAP in iso- and hypotonic state were upscaled to 400x400 pixels (Fiji’s bifurcation interpolation).

### Statistical analysis

Outlier detection was performed per condition × replicate group using two complementary methods. First, standard Tukey IQR fences were computed as Q1 − 1.5 × IQR and Q3 + 1.5 × IQR; data points falling outside these fences were flagged. Second, a modified z-score based on the median absolute deviation (MAD) was computed, with values exceeding a threshold of 3.5 flagged as outliers. Data points flagged by both methods were automatically excluded; those flagged by only one method were individually inspected and a manual retain or exclude decision was applied. Data points flagged by neither method were retained without further review.

Graphs were generated in GraphPad Prism 11 (11.0.0). Data was normalized across all conditions within each biological replicate prior to pooling as stated by min.-max. normalization or by transformation to mean of controls. Statistical analysis was performed with GraphPad Prism or in R. Statistical significance was determined as follows: for comparison of two groups by two-tailed Student’s t test, for comparison of multiple groups by ordinary one-way ANOVA with post hoc Dunnett’s or Tukey’ HSD comparison, or Kruskal-Wallis test with post hoc Dunn’s comparison, as appropriate from normality testing. Individual punctum-level measurements of endolysosomal LysoSensor intensity and per-organelle MAM size were subjected to linear mixed models (LMM) analysis with the formula: Value ∼ Condition + (1|Replicate/Image), fitted by restricted maximum likelihood using the lmerTest package (v3.2.0) in R (v4.5.2). Satterthwaite’s method was used for denominator degrees of freedom and Kenward-Roger correction for estimated marginal means (emmeans package, v2.0.1) and post-hoc comparisons were assessed with Tukey HSD correction. For both datasets, individual images were nested within biological replicates as a random effect. Due to the present number of biological replicates (n = 2–4 replicates per condition due to unbalanced factorial design), effect sizes (Cohen’s d) and estimated marginal means (EMMs) with 95% confidence intervals are reported rather than p-values alone. Distribution skewness (γ) was computed per condition and per replicate as a descriptive measure of subpopulation effects. The number of experimental replicates and the p-values can be found in the figure legends for each experiment.

### Data availability

The proteomic raw datasets generated for this study are available.

### Code availability

Custom Fiji macros (RS-FISH batch detection, FWHM-based ROI expansion), Python scripts (stacked line profiles, IQR and size-range-based outlier removal, colocalization analysis), and R scripts (LMM statistics) are available upon request.

---

**Figure S1.**
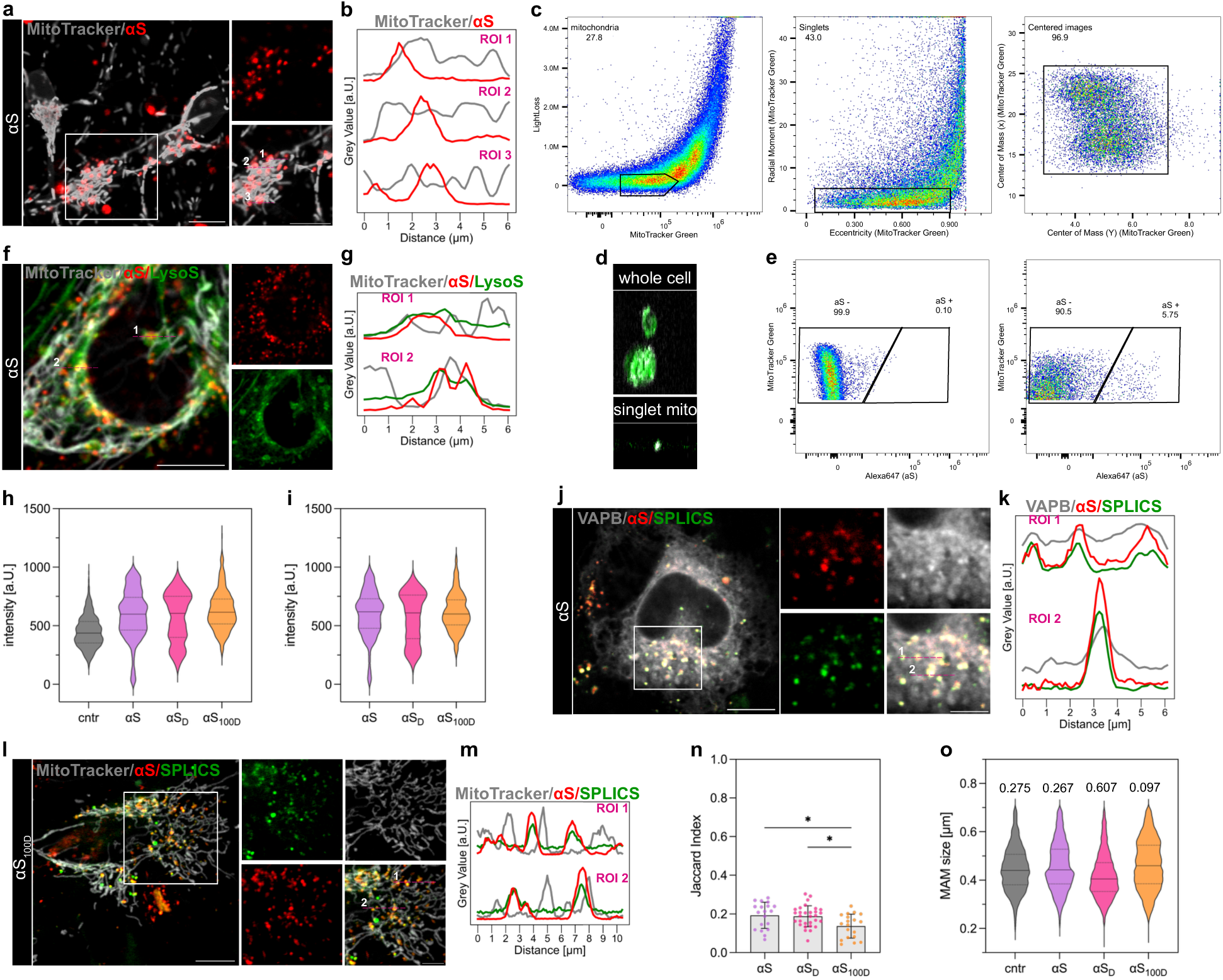
Subcellular distribution and compartmental characterization of internalized αS. a, Representative confocal images of dopaminergic neurons labeled with MitoTracker (grey) following treatment with αS (red), showing mitochondrial-associated αS puncta. Insets highlight regions of close spatial association. b, Intensity line profiles across selected ROIs from (a) demonstrating spatial proximity between mitochondria and αS puncta. c, Gating strategy for FAOS sorted mitochondria, combining spectral and BD CellView image-based features. Representative FAOS density plots showing sequential morphological gating of mitochondrial events: MitoTracker Green fluorescence versus axial light loss (P1), eccentricity versus radial moment to exclude doublets and aggregates (P2), and centre-of-mass gating to retain image-centred events (P3). Percentages indicate the proportion of events within each gate. d, Representative BD CellView images of sorted events confirm selection of morphologically small, circular mitochondrial entities. Top: intact cells present in the lysate prior to gating. Bottom: event classified as individual mitochondria (singlet mitochondria) based on MitoTracker Green fluorescence and morphological features, corresponding to the applied P1–P3 gate hierarchy. e, Representative FAOS density plots showing gating used to discriminate αS**^-^**/MitoTracker**^+^**from αS**^+^**/MitoTracker**^+^** fractions, based on spectral MitoTracker Green and Alexa647 fluorescence. Percentages indicate the proportion of events within each gate f, Confocal images of SH-SY5Y cells with MitoTracker, αS, and LysoSensor (green), revealing localization of αS puncta with endolysosomal compartments, in close apposition to mitochondria. Insets highlight regions containing mitochondria, αS and endolysosomes. g, Intensity line profiles from ROIs in (e) demonstrating overlap between αS and LysoSensor. h, i, Quantification of LysoSensor fluorescence intensity across endolysosomal compartments in neurons under control and αS-treated conditions. h, Total endolysosomal pool. For αS-treated conditions, this represents the sum of αS-associated and non-associated endolysosome populations. i, Intensity of αS-associated endolysosomes. LMM with nested random effects (1|Replicate/Image), Tukey HSD post-hoc, lmerTest. *N=*2-3 biological replicates; 6–9 FOV and 95–530 puncta per condition, per ROI. j, Confocal images of SH-SY5Y expressing SPLICS (green) and VAPB (grey) and conditioned with αS (red). Insets highlight regions of colocalization. k, Intensity line profiles from ROIs in (j) confirming colocalization of αS with VAPB- and SPLICS-positive structures. l, Representative confocal images of SH-SY5Y with MitoTracker, αS_100D_, and expressing SPLICS (green), confirming that αS_100D_ shows comparable localization to mitochondria tubules and MAM-puncta as observed for αS and αS_D_. Insets show magnified views of contact regions. m, Intensity line profiles from ROIs in (l) demonstrating overlap between αS_100D_ and SPLICS-positive contact sites. n, Jaccard index of object-based colocalization between αS and SPLICS, providing a symmetric, size-independent measure of spatial overlap. Ordinary one-way-Anova with post hoc Tukey’s. *N=*2-3 biological replicates. o, MAM contact site radius (µm) across conditions. No significant difference in mean MAM size detected. Distribution analysis revealed condition-dependent skewness (γ): αS_D_ showed a 2-fold increase in skewness factor relative to control (αS_D_ γ = 0.603 vs ctrl γ = 0.275), consistent across both biological replicates (0.606, 0.581), suggesting a subpopulation shift toward smaller contact sites, while αS_100D_ showed the lowest γ (0.096). LMM with nested random effects (1|Replicate/Image), lmerTest; *N=*2 biological replicates. Skewness: per-replicate γ computed on image-mean-residualized values (scipy.stats.skew). Bar plots show mean ± s.e.m: each point represents a single FOV. * p < 0.05. Scale bars: a, e, i, m, 10 µm; insets i, m, 5 µm. Violin plots display the distribution of individual data points, with the median and interquartile range.

**Figure S2.**
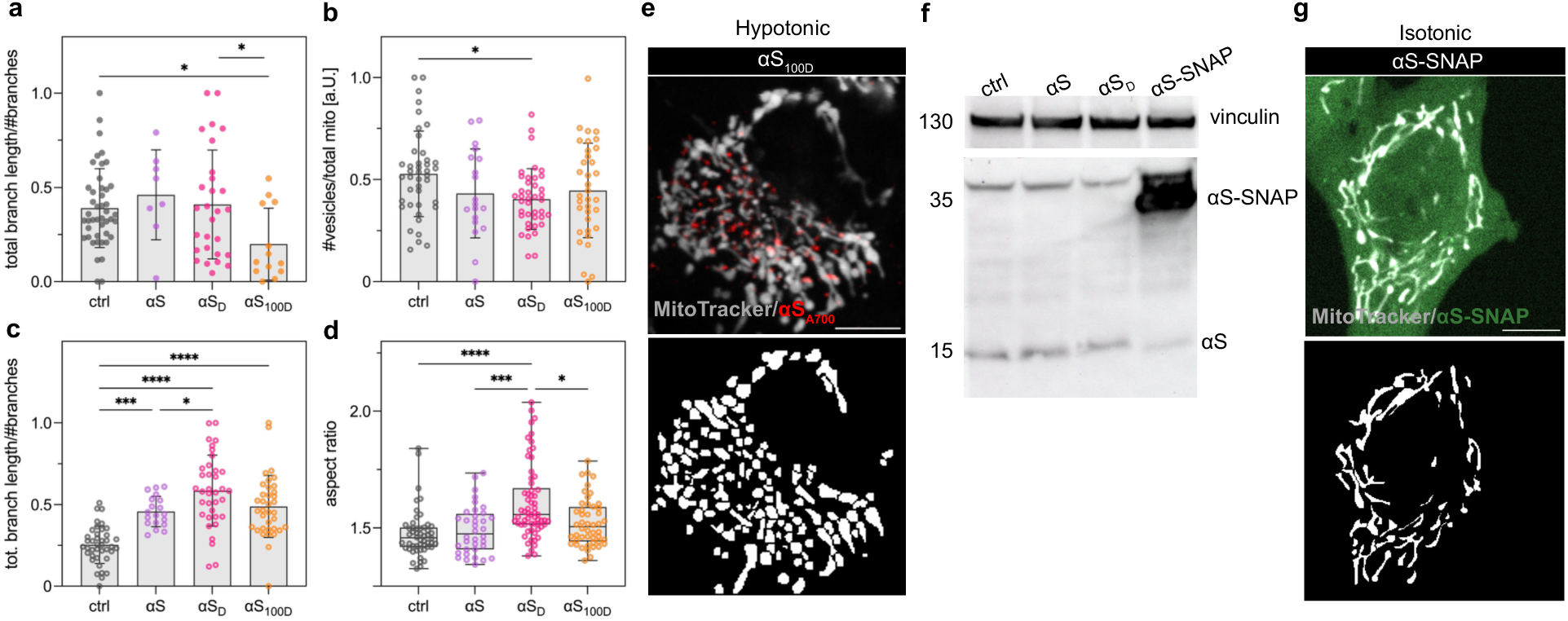
Spreading αS constrains mitochondrial remodeling in a C-terminal-dependent manner and independently of intracellular αS expression. a, Quantification of mitochondrial mean branch length under homeostatic (isotonic) conditions across conditions. αS does not show significant alterations, while αS_100D_ shows reduced branch length under homeostatic conditions. Ordinary one-way-Anova with Dunnett’s post-hoc. *N=2-3* biological replicates. b-d, Quantification of mitochondria morphology post-hypotonic treatment in control and αS-treated dopaminergic neurons. The number of LICVs shows a significant reduction in the αS_D_ condition (b), while mean branch length (c) and branch elongation (d) are significantly increased in a concentration- and C-terminal-dependent manner. Ordinary one-way-Anova with Tukey’s or Kruskal-Wallis with Dunn’s post-hoc. *N=2* biological replicates. e, Representative image and segmentation mask of mitochondria in αS_100D_-treated SH-SY5Y cells under hypotonic conditions. f, Western blot of cell lysates from ctrl, αS-treated (αS, αS_D_), and αS-SNAP-overexpressing SH-SY5Y cells, probed with anti-αS and anti-SNAP antibodies. Vinculin was used as a loading control. g, Representative confocal image and binary mask of αS-SNAP expressing SH-SY5Y cells under isotonic conditions. Data are presented as mean ± s.e.m.; each point represents a single FOV. * p < 0.05, ** p < 0.01, *** p < 0.001, **** p < 0.0001. Filled markers: isotonic; open markers: hypotonic. Scale bars: 10 µm.

**Figure S3.**
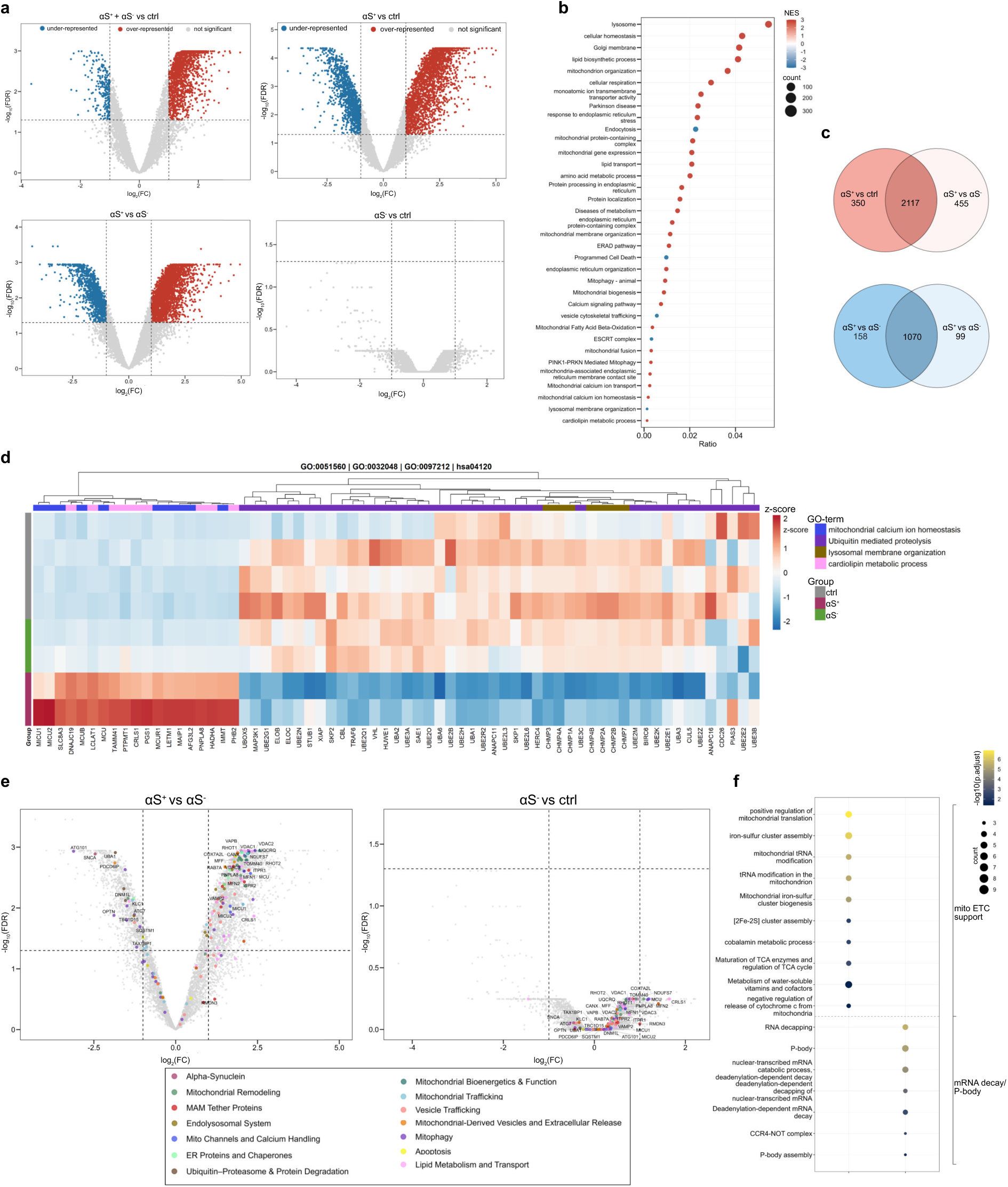
Differential proteomic changes and pathway stratification across αS–defined mitochondrial populations. a, Volcano plots showing differentially abundant proteins across all pairwise comparisons: combined αS-treated mitochondrial populations (αS⁺ + αS⁻) vs ctrl, αS⁺ vs ctrl, αS⁺ vs αS⁻, αS⁻ vs ctrl. Proteins are colored to indicate over-representation (red), under-representation (blue), or no significance (gray). Volcanoes illustrate global proteomic changes induced by αS exposure, which is absent in αS⁻ vs ctrl. b, GSEA of pooled mitochondrial populations from αS-treated cells (αS⁺ + αS⁻) vs ctrl. Enriched pathways include mitochondrial organization, respiration, lipid metabolism, and ER stress, whereas trafficking and transport terms are depleted. c, Venn diagrams showing overlap of differentially expressed proteins across comparisons. Top: overlap of overrepresented proteins. Bottom: overlap of underrepresented proteins. d, Heatmap of selected enriched pathways across ctrl, αS⁻, and αS⁺. Hierarchical clustering reveals distinct pathway signatures, with αS⁺ mitochondria enriched in mitochondrial calcium homeostasis and lipid metabolism and depleted in ubiquitin-mediated processes. d, ORA of unique pathways distinguishing αS⁺ vs ctrl and αS⁺ vs αS⁻ comparisons. Dot plot highlights pathways selectively enriched or depleted in αS⁻ vs ctrl. e, Volcano plots for: top, αS⁺ vs ctrl; middle, αS⁺ vs αS⁻; bottom, αS⁻ vs ctrl. These comparisons illustrate the magnitude of proteomic changes in αS⁺ mitochondria relative to both control and αS⁻, and the comparatively limited changes in αS⁻ vs ctrl. f, Volcano plots comparing αS⁺ vs αS⁻ (left) and αS⁻ vs ctrl (right) mitochondrial fractions. Proteins of interest are highlighted and grouped into custom-defined functional categories, whereas all other differentially abundant proteins are shown in grey. Annotated proteins are referenced in the Results section. Dashed lines indicate significance thresholds (adjusted p < 0.05; |log₂FC| > 1). A table listing proteins of interest and their assigned functional categories is provided below (Supplementary Table S1). Statistical significance was determined using the false discovery rate (FDR) correction, further described in the Methods. Biological replicates, ctrl, αS, *N=*4; αS_D_, *N=*2.

**Figure S4.**
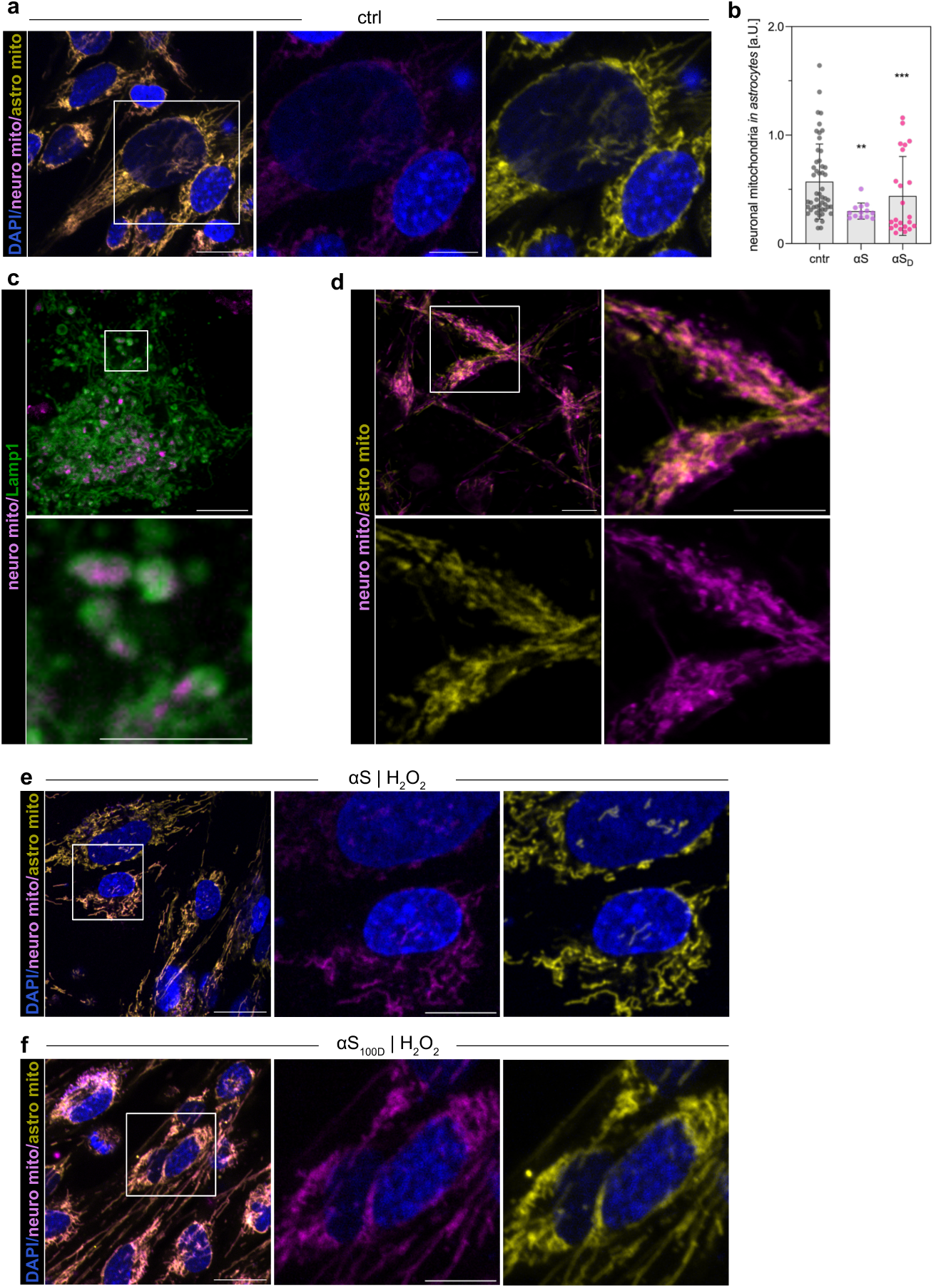
αS impairs neuronal mitochondrial transfer and lysosomal processing. a, Representative confocal images of astrocytes co-cultured with neurons under basal conditions, showing detectable uptake of neuronal mitochondria (denoted neuro mito: magenta) into astrocytes (denoted astro mito: yellow). Insets highlight transferred mitochondria content. b, Quantification of a, neuronal mitochondrial transfer to astrocytes under basal conditions. αS presence in neurons significantly reduces mitochondrial transfer to astrocytes compared with controls. LMM with random intercept (1|Replicate), Tukey HSD post-hoc on estimated marginal means, lmerTest/emmeans. *N=*2 biological replicates. c, High-resolution images showing neuronal mitochondria (magenta) localized within Lamp1-positive lysosomal compartments (green) in astrocytes, indicating delivery of transferred neuronal mitochondria to degradative organelles. d, Representative images of the neuronal portion of co-cultures. Images suggest the incorporation of astrocytic mitochondrial signal (yellow) into the neuronal mitochondrial network (magenta). e, f, Representative images of astrocytes co-cultured with αS and αS_100D_-treated neurons under oxidative stress. Neuronal mitochondrial signal (magenta) illustrates neuron-to-astrocyte mitochondrial transfer. Data are presented as mean ± s.e.m.; each point represents an individual cell or field of view. * p < 0.05, ** p < 0.01, *** p < 0.001. Values normalized to within-replicate ctrl+H₂O₂ mean. Filled markers: no insult; open markers: +H₂O₂. Scale bars: a, c, e, f 20 µm; d, insets a, e, f 10 µm, inset c, 2.5 µm.

